# A Cytokine Receptor Signaling Atlas Reveals How STAT Mosaics Fine-Tune T Cell Function

**DOI:** 10.64898/2026.08.19.745852

**Authors:** Pingdong Tao, Ruchir Rastogi, Hua Jiang, Yang Zhao, Leon L. Su, Kevin Jude, Anshul Kundaje, K. Christopher Garcia

## Abstract

The extent to which JAK/STAT cytokine signaling is functionally redundant or selective remains debated. Here we engineered a “double” orthogonal IL-2/IL-2Rβ/γc ternary system enabling programmable, interference-free activation of each of the 36 mammalian cytokine receptors, and their downstream six STATs, in T cells. At the membrane-proximal level, comprehensive phospho-signaling profiling revealed that while each receptor activates a dominant STAT, unique STAT activation fingerprints derived from combinatorial biases fine-tune nuanced T cell fates. At the membrane-distal level, single-cell transcriptomic atlas of all cytokine receptors confirmed that these STAT mosaics sensitively specify non-redundant transcriptional programs. STAT5-dominant receptors drove proliferative expansion at the expense of stemness; STAT3-driven programs instructed a continuum from stem cell memory to terminal effector states with preserved cytotoxic capacity and mediated superior curative antitumor responses; while other STATs specified highly restricted phenotypes. These findings decode a STAT signaling vocabulary that defines the intrinsic functional bandwidth of natural cytokines.

## Introduction

Cytokine receptors are activated by ligand-induced homo- or heterodimerization, leading to transphosphorylation of receptor-associated JAK kinases and subsequent activation of STAT transcription factors that instruct diverse gene expression programs underlying immune cell development and differentiation^1–5^. Despite sharing a relatively limited set of six STAT family members, the 36 known cytokine receptors collectively encode highly diverse and context-dependent cellular outcomes^6–10^. This raises a fundamental question: How does a limited STAT vocabulary encode the full functional complexity of different cytokine receptors? Decades of study and extensive transcriptomic profiling have characterized cytokine-induced responses across immune cell types^11–14^. Cytokine receptors generally activate a dominant STAT (i.e. IL-2 is a STAT5 cytokine), however the degree to which the other five STATs, cooperate to fine-tune cytokine-specific versus pleiotropic functions remains incompletely understood, particularly in T cell subsets^15–17^.

This question has also become increasingly important for developing T cell-based immunotherapies. T cells supplemented with immunomodulatory cytokines have been widely employed to enhance antitumor efficacy in adoptive cell therapy^18–21^. Synthetic biology approaches enforce expression of cytokine receptors to drive JAK/STAT signaling for adoptive cell therapy (ACT), aiming to overcome the restricted expression patterns of endogenous receptors on T cells and to harness the untapped signaling diversity enabled by ectopic receptor engineering^22–27^. Defined STAT-recruiting motifs have similarly been incorporated into synthetic receptor platforms to modulate T cell function^28–30^. However, these advances have largely proceeded empirically, reflecting a fundamental lack of understanding of how specific STAT signaling combinations encode distinct cellular programs. Resolving this question would provide a mechanistic framework for decoding cytokine signaling language and a blueprint for the rational design of engineered cell therapies.

A limitation of prior experimental systems to study this question was the lack of clear insulation of cytokine signaling pathways from background interference by co-mingling endogenous signaling receptors and cytokine. In an attempt to overcome this limitation, we previously engineered a “single” orthogonal cytokine receptor system based on the IL-2 and IL-2Rβ site 1 interaction^31,32^. By pairing the engineered ortho-IL-2Rβ with endogenous common γc, we could selectively activate authentic IL-2 signaling on the target cell using an orthogonal IL-2. We extended this orthogonal approach to manipulate the activation of more cytokine receptors including γc family receptors^33^ and non-γc family receptors^34^, and found that distinct cytokine receptor pairings generated signals that, unexpectedly, directed the engineered T cells toward a broader phenotypic spectrum. Yet, these phenomenological observations highlighted that a complete and reductionist understanding of cytokine receptor-mediated functional specificity and redundancy, at both membrane-proximal and distal levels, remains lacking. Given that cytokine receptors activate multiple signaling pathways, including canonical JAK-STAT as well as MAPK and PI3K-mTOR cascades^35–37^, how these pathways are integrated to specify T cell functional programs also remains unclear.

Here we developed a fully orthogonal cytokine-receptor system, giving us complete synthetic control over both chains of the receptor dimer that initiates JAK/STAT and non-STAT pathways. Using this platform we generated phospho-STAT maps for all cytokine receptor intracellular domains. In parallel, we used single-cell RNA sequencing to generate a comprehensive transcriptional atlas across the same orthogonal cytokine receptor conditions in mouse CD8^+^ T cells, and reconstructed differentiation trajectories associated with STAT-dependent transcriptional programs. Through systematic *in vivo* profiling and evaluation of receptors preferentially enriched among tumor-infiltrating T cells, we find that distinct, finely-tuned STAT activation fingerprints specify profoundly different T cell states and antitumor outcomes, with STAT3-dominant signaling conferring uniquely superior curative efficacy. Our results build on previous studies of STAT hierarchies^9,38,39^, and suggest a model whereby STAT combinatorial biases, not a single dominant STAT, act as the determinants of selective T cell phenotype and function, establishing the boundary conditions for the functional space accessible to both natural and engineered receptors.

## Results

### A double-orthogonal cytokine receptor system based on IL-2/IL-2Rβ/γc complex

Although our previously engineered single-orthogonal IL-2/IL-2Rβ system enabled selective cytokine signaling, it remained dependent on endogenous γc, limiting complete insulation from native cytokine receptor networks. To overcome this limitation and establish a fully reductionist synthetic system to isolate signaling by each receptor intracellular domain, we engineered a double-orthogonal IL-2 receptor system composed of ortho-IL-2, ortho-IL-2Rβ, and ortho-γc (**Fig. 1A**), in which signaling requires the coordinated assembly of two orthogonal receptor chains, enabling selective activation of engineered cells without engaging endogenous cytokine receptors.

**Figure 1.**
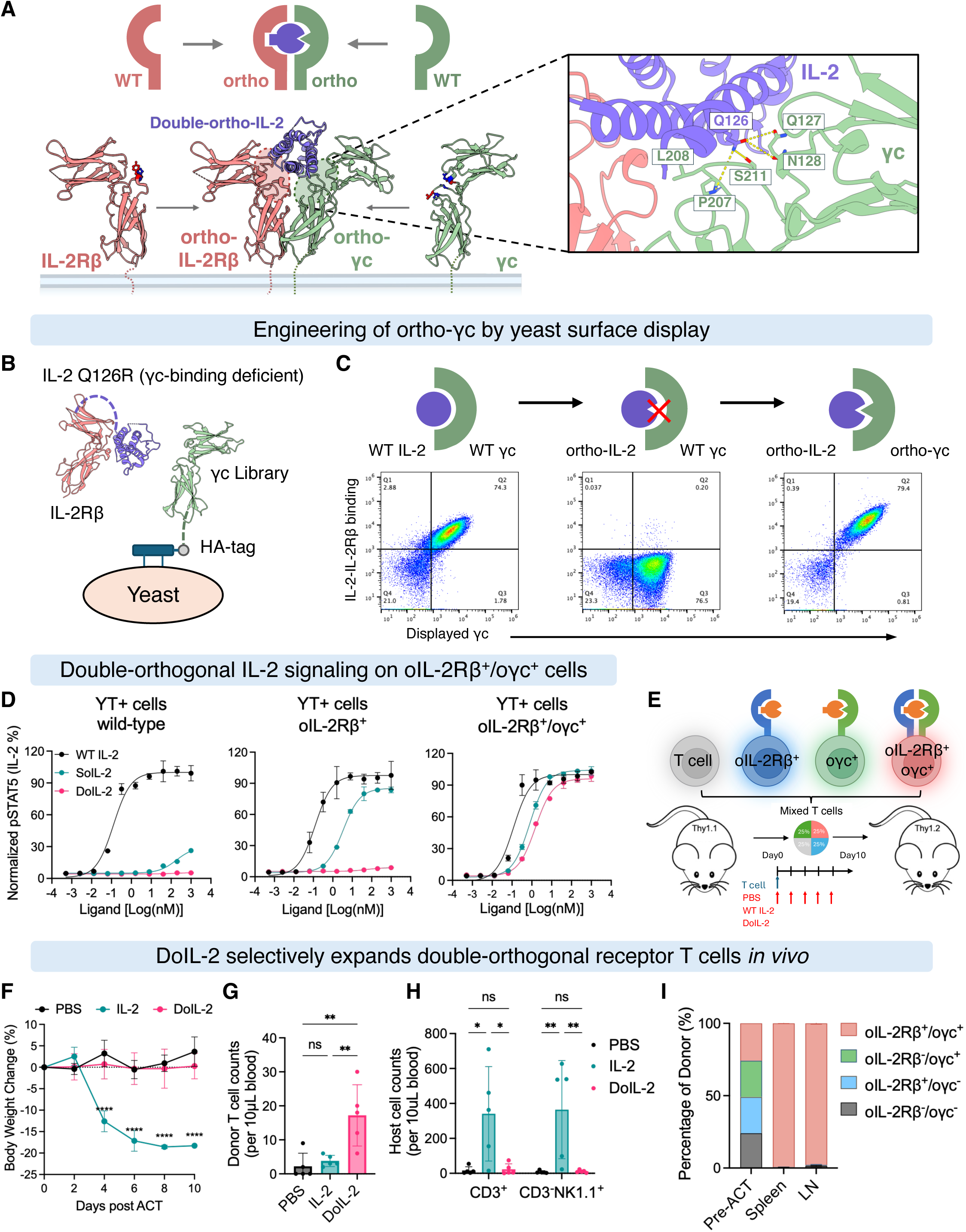
A double-orthogonal IL-2/receptor system enables selective, interference-free activation of engineered T cells *in vivo*. (A) Schematic of the double-orthogonal ternary signaling complex. A mutant IL-2 (DoIL-2) is engineered to bind exclusively to orthogonal IL-2Rβ (ortho-IL-2Rβ) and orthogonal γc (ortho-γc) and cannot engage their wild-type counterparts on normal immune cells. Structural basis for orthogonal γc engineering. Key γc residues (Q127, N128, P207, L208, S211) that contact IL-2 (Q126) in the natural complex were targeted for mutagenesis, based on the human IL-2R crystal structure (PDB: 2b5i). (B) Strategy for engineering orthogonal ãc by yeast surface display. An IL-2 mutein fused to the IL-2Râ extracellular domain was used to select for ãc variants that bind mutant but not wild-type IL-2. (C) Binding validation of selected ortho-ãc candidates on yeast surface. Wild-type ãc binds wild-type IL-2-IL-2Râ fusion (left) but not the Q126R mutant (middle); ortho-ãc binds the Q126R mutant (right). (D) STAT5 signaling validation of engineered orthogonal IL-2/receptor system using YT+ cells. Wild-type cells responded only to wild-type IL-2. Cells expressing ortho-IL-2Rβ responded to single-orthogonal IL-2 (SoIL-2) but not double-orthogonal IL-2 (DoIL-2), whereas only cells co-expressing ortho-IL-2Rβ and ortho-γc responded to DoIL-2. (E) Schematic of *in vivo* orthogonality validation. Mice received co-transfer of T cells expressing ortho-IL-2Rβ alone (TagBFP^+^), ortho-γc alone (ZsGreen^+^), or both receptors (double-orthogonal; mCherry^+^), followed by treatment with PBS, IL-2, or DoIL-2. (F-I) Selectivity of engineered T cell expansion *in vivo*. Systemic toxicity assessment by body weight change (F). Quantification of donor T cells (G) and host T and NK cells (H) in peripheral blood 5 days after treatment. DoIL-2 selectively expands double-orthogonal donor T cells without activating host lymphocytes. Quantification of the proportion of donor T cells expressing the double-orthogonal receptor (mCherry⁺) in the spleen and lymph nodes 10 days after DoIL-2 treatment (I). See also **Figure S1** and **S2**.

To engineer the orthogonal γc interface, structural analysis identified Q126 within helix D of human IL-2 as a conserved residue critical for γc engagement across the γc cytokine family (**Fig. 1A**)^40,41^. Saturation mutagenesis at this position identified several charge-altering variants (Q126X) with disrupted γc binding, which were selected as candidate double-orthogonal IL-2 molecules (**Fig. S1A**). We then evolved compensatory γc variants using a yeast-display platform with an IL-2/IL-2Rβ fusion protein to stabilize γc binding (**Fig. 1B**, **Fig. S1B-D**). Following four rounds of selection (**Fig. S1E-F**), clone R1 was identified as the optimal orthogonal γc, selectively restoring robust binding to the IL-2 Q126R mutein (**Fig. 1C**). We therefore designated the single ortho-IL-2 (1A1, SoIL-2)^32^ with Q126R mutation as double ortho-IL-2 (DoIL-2) and the R1 variant as ortho-γc.

To evaluate signaling specificity, we expressed orthogonal IL-2Rβ and/or orthogonal γc in YT+ reporter cells and primary human T cells. DoIL-2 failed to induce signaling in untransduced cells or cells expressing either receptor chain alone, but robustly activated STAT5 in cells coexpressing both orthogonal receptors, reaching levels comparable to wild-type IL-2 (**Fig. 1D**, **Fig. S2A-C**). To assess *in vivo* orthogonality, we generated chimeric orthogonal receptors containing engineered human extracellular domains fused to mouse transmembrane and intracellular signaling domains (**Fig. S2D**). DoIL-2 selectively activated STAT5 signaling only in T cells coexpressing both orthogonal receptor chains (**Fig. S2E**). Following adoptive transfer of mixed single-or double-orthogonal receptors engineered T cells (**Fig. 1E**), treatment with MSA-DoIL-2 selectively expanded double-orthogonal receptor-expressing donor T cells without detectable systemic toxicity, whereas wild-type IL-2 induced substantial toxicity and broadly expanded endogenous lymphocyte populations (**Fig. 1F-I**).

Leveraging the conserved architecture of the IL-2 receptor complex, we further generated a fully murine double-orthogonal platform using the same engineering principles for *in vivo* use in order to bypass immunogenicity of the human receptors. Mouse DoIL-2 selectively activated STAT5 only in cells coexpressing both orthogonal receptor chains, albeit with a higher EC50 relative to the human system (**Fig. S2F**).

In summary, we established a fully orthogonal IL-2/IL-2Rβ/γc signaling platform that enables selective activation and expansion of engineered cells without engaging endogenous cytokine receptor networks.

### Isolating individual cytokine receptor signaling using the double-orthogonal system

To evaluate whether the double orthogonal system could recapitulate signaling across the full cytokine receptor repertoire, we enumerated the 31 naturally occurring dimeric cytokine receptor pairs (i.e. homo- and heterodimeric receptors) in mouse and generated all 36 unique orthogonal receptor chimeras by replacing their extracellular domains with ortho-IL-2Rβ or ortho-γc. We adopted the human-mouse chimeric orthogonal receptor configuration throughout, as human DoIL-2 exhibited superior affinity and signaling potency. Phospho-STAT profiling in primary mouse T cells expressing orthogonal receptor pairs demonstrated that DoIL-2 robustly activated all engineered receptor dimers under naturally occurring pairings (**Fig. 2A-B, Fig. S3A-B**).

**Figure 2.**
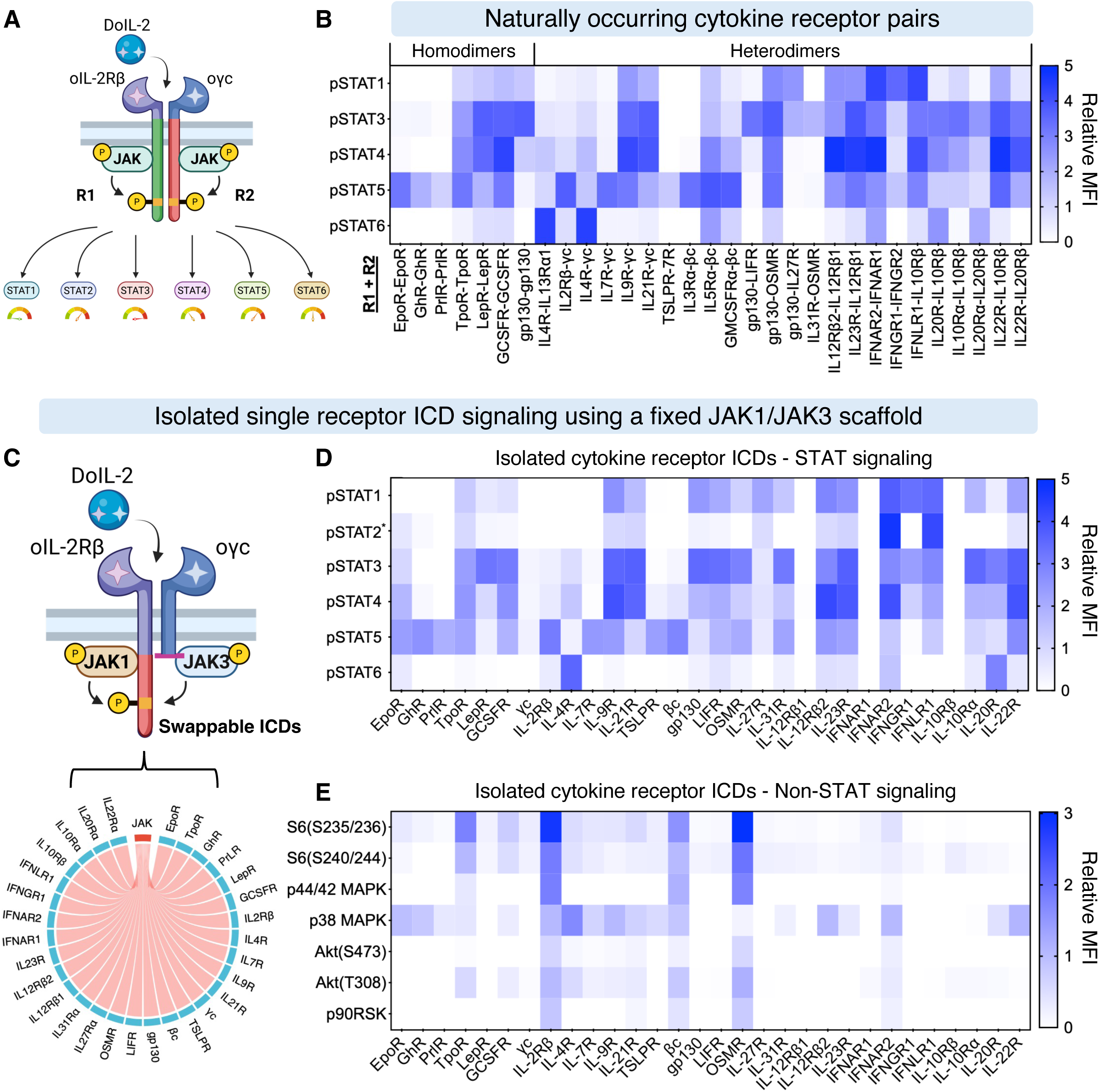
Every JAK-associated cytokine receptor encodes a quantitatively distinct STAT activation fingerprint. (A) Schematic of the natural receptor pair reconstruction approach. The intracellular domains (ICDs) of both chains in naturally occurring cytokine receptor pairs were swapped into the double-orthogonal system, allowing each natural pair to be selectively activated by DoIL-2 without interference from endogenous receptors. (B) Heatmap of pSTAT1/3/4/5/6 activity across all 31 reconstructed natural receptor pairs. Each receptor pair generates a characteristic STAT activation pattern, quantified as log2 fold change (log2FC) over Box control by phospho-flow in mouse T cells stimulated with DoIL-2 (100 nM). Data collected in duplicate. (C) Schematic of the decoupled receptor approach for isolating individual ICD contributions. A chimeric ortho-IL-2Râ receptor carries the fixed JAK-binding region and the ICD of interest; a truncated ortho-ãc contributes only JAK recruitment. This allows the STAT-recruiting activity of any single receptor ICD to be measured in isolation. (D-E) Heatmaps of pSTAT1/2/3/4/5/6 (D) and selected non-STAT phospho-signaling (E) activities across all 30 decoupled receptor ICDs profiled using the approach in (C). Values are log2FC over Box control. See also **Figure S3** and **S4**.

While the pSTAT profiles of natural receptor dimeric assemblies are informative, they reflect the combined output of two receptor chains signaling together, confounding our desire to reduce the signaling output to a single receptor. To isolate the intrinsic signaling contribution of each individual receptor intracellular domain (ICD), we redesigned the system to read out a single ICD at a time. The strategy, illustrated in **Fig. 2C**, preserves the orthogonal extracellular and transmembrane regions of the receptor scaffold but strips signaling capacity from the ortho-γc chain, retaining only the membrane-proximal Box region required for JAK3 recruitment^42^, with no remaining intracellular tail that could contribute STAT-recruiting activity. The ICD of any “guest” receptor is then fused to the ortho-IL-2Rβ chain. In this configuration, DoIL-2 brings the two chains together, but all downstream STAT signaling is directed exclusively through the single test ICD on the IL-2Rβ arm, reducing a two-chain signaling complex to a single-ICD readout.

An initial version of this design, in which the transmembrane and membrane-proximal Box regions of IL-2Rβ were left as native sequence, failed to produce robust STAT phosphorylation for a subset of receptors (**Fig. S3C-E**). We resolved this by fixing both the transmembrane domain and the membrane-proximal Box regions^43,44^ of IL-2Rβ as invariant scaffolding, creating a uniform IL-2Rβ-JAK1/γc-JAK3 signaling platform that ensures consistent transphosphorylation of every test ICD regardless of its origin. This JAK1/JAK3 pairing is particularly well suited as a universal scaffold because JAK1 and JAK3 have evolved for signaling promiscuity through their central role in the γc cytokine receptor hub. The result is a system that reports the intrinsic activity of the minimal functional unit of a cytokine receptor: the ICD segment downstream of the Box1/Box2 JAK-binding motif, which contains the adaptor and STAT docking sites^38^.

Applying this optimized platform to 30 receptors (excluding 6 with minimal intracellular domains) revealed robust STAT activation across the majority of tested ICDs, with the exception of a subset of short-tailed receptors with no apparent STAT binding sites, γc, IFNAR1, IL-12Rβ1, and IL-10Rβ, which remained minimally responsive across all STATs (**Fig. 2D, Fig. S4A-B**).

Across the STAT family, STAT1, STAT3, STAT4, and STAT5 displayed broad activation across receptors, whereas STAT2 and STAT6 exhibited highly restricted patterns. STAT2 activation was confined to type I and type III interferon receptors (IFNAR2 and IFNLR1), while STAT6 activation was largely restricted to IL-4R, with modest activation observed in IL-20R and IFNAR2. Notably, most receptors exhibited combinatorial activation of STAT1/3/4/5 with distinct signaling biases. For example, both IL-12Rβ2 and IL-23R activated multiple STATs, but IL-23R preferentially induced STAT3, whereas IL-12Rβ2 favored STAT4. Similarly, IL-9R and IL-21R showed comparable STAT1/3/4 activation but differed in STAT5 engagement.

In addition to canonical STAT signaling, we extended the phosphorylation profiling to selected non-STAT pathways that also contribute to the downstream signaling network, including MAPK (ERK1/2, p38), PI3K (Akt), and ribosomal signaling (S6 and S6K). In contrast to the broadly distributed STAT responses, robust activation of these pathways was restricted to specific receptors, including TpoR, GCSFR, IL-2Rβ, βc, and OSMR, although p38 MAPK activation showed broader coverage (**Fig. 2E, Fig. S4C**).

Taken together, the double orthogonal system enables receptor-by-receptor dissection of cytokine signaling free from native pairing. Each receptor generates a quantitatively distinct STAT activation fingerprint defined by receptor-specific stoichiometries across STAT and non-STAT pathways.

### Transcriptomic atlas of T cells across all cytokine receptor conditions

Having shown that each cytokine receptor encodes a distinct quantitative phospho-STAT signaling fingerprint, we next asked whether these signaling programs specify distinct T cell states by transcriptionally profiling cells activated through each receptor. We performed high-throughput single-cell transcriptomic profiling across all the same activated orthogonal receptors in mouse CD8⁺ T cells that we generated pSTAT activation profiles. Pmel-1 T cells transduced with double-orthogonal receptors or control were stimulated with DoIL-2 for 24 hours, followed by sequencing using a split-pool combinatorial barcoding platform (**Fig. 3A**). In total, 35 samples were profiled, including 30 natural receptors, 3 engineered receptors, and 2 controls: a non-transduced (NT) control and a Box mock receptor containing only the JAK1/JAK3-binding region (no STAT binding sequences). After quality filtering, 63,669 cells were retained for downstream analysis, with per-sample cell numbers ranging from 1,101 to 2,370. These cells were integrated across samples, and the resulting UMAP embedding revealed a comprehensive transcriptomic atlas of CD8⁺ T cells driven by each cytokine receptor (**Fig. 3B**).

**Figure 3.**
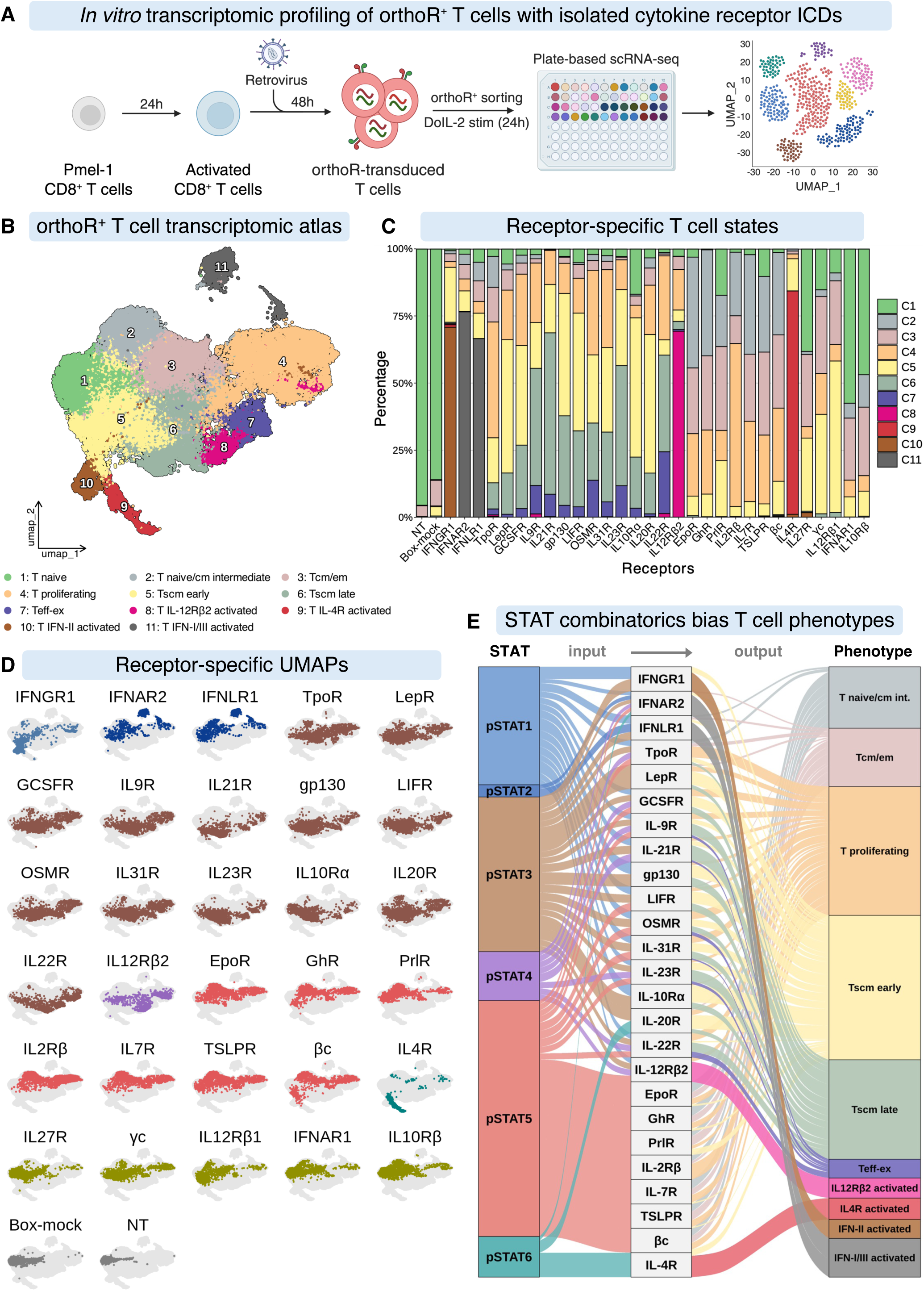
Distinct STAT signaling fingerprints deterministically specify distinct T cell transcriptional states. (A) Experimental workflow for *in vitro* single-cell transcriptomic profiling. Pmel-1 CD8^+^ T cells were transduced with each of 30 orthogonal receptor constructs, sorted for receptor expression, stimulated with DoIL-2 (100 nM, 24 h), and processed for scRNA-seq using a plate-based combinatorial barcoding platform. (B) UMAP of 58,624 Pmel-1 CD8^+^ T cells from all receptor conditions. Eleven transcriptionally distinct clusters were identified and annotated based on marker gene expression. (C) Stacked bar plot showing the proportion of cells per annotated cluster for each receptor condition, illustrating how each receptor drives cells into distinct cluster distributions. (D) Per-condition UMAPs showing T cell distributions across all 30 receptor conditions and two controls, colored by the dominant STAT signaling category assigned to each receptor. Conditions with similar STAT profiles cluster together in transcriptional space. (E) Sankey plot linking quantitative STAT activation profiles (from Figure 2D) to T cell cluster distributions (from Figure 3C), revealing a direct correspondence between receptor signaling input and transcriptional output. See also **Figure S5**.

Cells resolved into 11 distinct clusters representing a spectrum of recognizable T cell states (**Fig. S5A-B**). Seven of these correspond to canonical states found in normal immune responses: a naïve-like state (C1); a naïve/central memory intermediate (C2); central and effector memory T cells (C3); actively proliferating T cells (C4); an early stem cell memory state (C5, or early Tscm); a more differentiated late stem cell memory state (C6, late Tscm) with early signs of effector gene expression (*Gzmb*, *Ifng*); and an exhausted-like effector state (C7) that carries both cytotoxic and inhibitory receptor markers. Four additional clusters were almost entirely occupied by cells from a single receptor condition, indicating that certain receptors drive a transcriptional program so distinctive it has no equivalent anywhere else in the atlas: a cluster specific to IL-12Rβ2 (C8), one specific to IL-4R (C9), one specific to IFNGR1 (C10), and one shared by IFNAR2 and IFNLR1 (C11, both type I/III interferon receptors) (**Fig. S5C-D**).

Although 11 clusters defined this landscape across the total T cell population, cluster distribution and relative abundance varied substantially among receptor conditions (**Fig. 3C**). The Box control showed a distribution similar to the NT control, with both primarily localized within the undifferentiated T naïve cluster (C1). Every other receptor drove cells into more differentiated regions of the map, but the specific destinations differed by STAT program. Excluding the four receptor-specific clusters, STAT5-dominant receptors (EpoR, GhR, PrlR, IL-2Rβ, IL-7R, TSLPR, and βc) were predominantly enriched in T naïve /cm intermediate (C2) and Tcm/em (C3) clusters. This observation is consistent with the established roles of IL-7 and IL-15 in supporting naïve and memory T cell maintenance^45,46^. In contrast, receptors engaging multiple STAT1/3/4/5 signaling preferentially drove T cell differentiation toward Tscm-associated clusters (C5 and C6). Within this group, TpoR, GCSFR, gp130, LIFR, OSMR, and IL-31R exhibited a balanced distribution between early and late Tscm states. By comparison, LepR, IL-10R, and IL-20R were biased toward the early Tscm cluster (C5), whereas IL-9R, IL-21R, IL-22R, and IL-23R promoted further differentiation toward a more effector-poised late Tscm state (C6). Notably, receptors that supported Tscm differentiation also contributed to the generation of the exhausted-like effector subset (C7), suggesting a shared differentiation trajectory. Unexpectedly, several receptors with minimal STAT activation, including γc, IFNAR1, IL-12Rβ1, IL-10Rβ, and IL-27R, still elicited substantial transcriptional responses, in contrast to the Box control, indicating that these receptors are not functionally silent despite weak canonical STAT signaling (**Fig. 3C-D**). Collectively, receptor-specific UMAP distributions revealed that T cell state occupancy is closely associated with the dominant STAT signaling engaged by each receptor (**Fig. 3D, Fig. S5E-F**).

This transcriptomic atlas establishes a direct linkage between receptor-encoded membrane proximal signaling and T cell state. Receptors with similar quantitative STAT biases preferentially populated neighboring regions of the differentiation landscape while maintaining distinct transcriptional identities, demonstrating that quantitative differences in cytokine receptor signaling organize T cell differentiation into a structured continuum rather than discrete signaling modules. To present the full picture of how STAT signaling maps onto T cell fate in a single summary, we constructed a flow diagram connecting each receptor’s STAT activation profile to the T cell cluster distribution it produced in the atlas (**Fig. 3E**). This diagram shows that T cell phenotypic fragmentation occurs because of the receptor-specific mosaic of STAT activities. These data establish a deterministic signaling-to-state framework that defines how quantitative STAT combinatorics specify distinct T cell fates.

### STAT-dependent T cell differentiation trajectories

To understand how these fingerprints influence T cell differentiation, we scored each atlas cell across all receptor conditions for six T cell functional signatures: stemness, activation, effector molecules, exhaustion-associated markers, interferon-stimulated gene (ISG) responses, and proliferation (**Fig. S6A**), and reconstructed differentiation trajectories. This revealed that activation, effector, and exhaustion-associated signatures largely overlapped and were mutually exclusive with stemness, suggesting progressive differentiation from naïve-like cells toward distinct functional states, whereas ISG and proliferation signatures remained largely confined to discrete populations (**Fig. 4A, Fig. S6B-F**). Restricting trajectory analysis to non-cycling T cells and inferred pseudotemporal lineages using a cluster-based lineage inference approach^47^. This analysis identified six differentiation trajectories that closely aligned with receptor-specific dominant STAT signaling programs (**Fig. 4B, Fig. S6G**).

**Figure 4.**
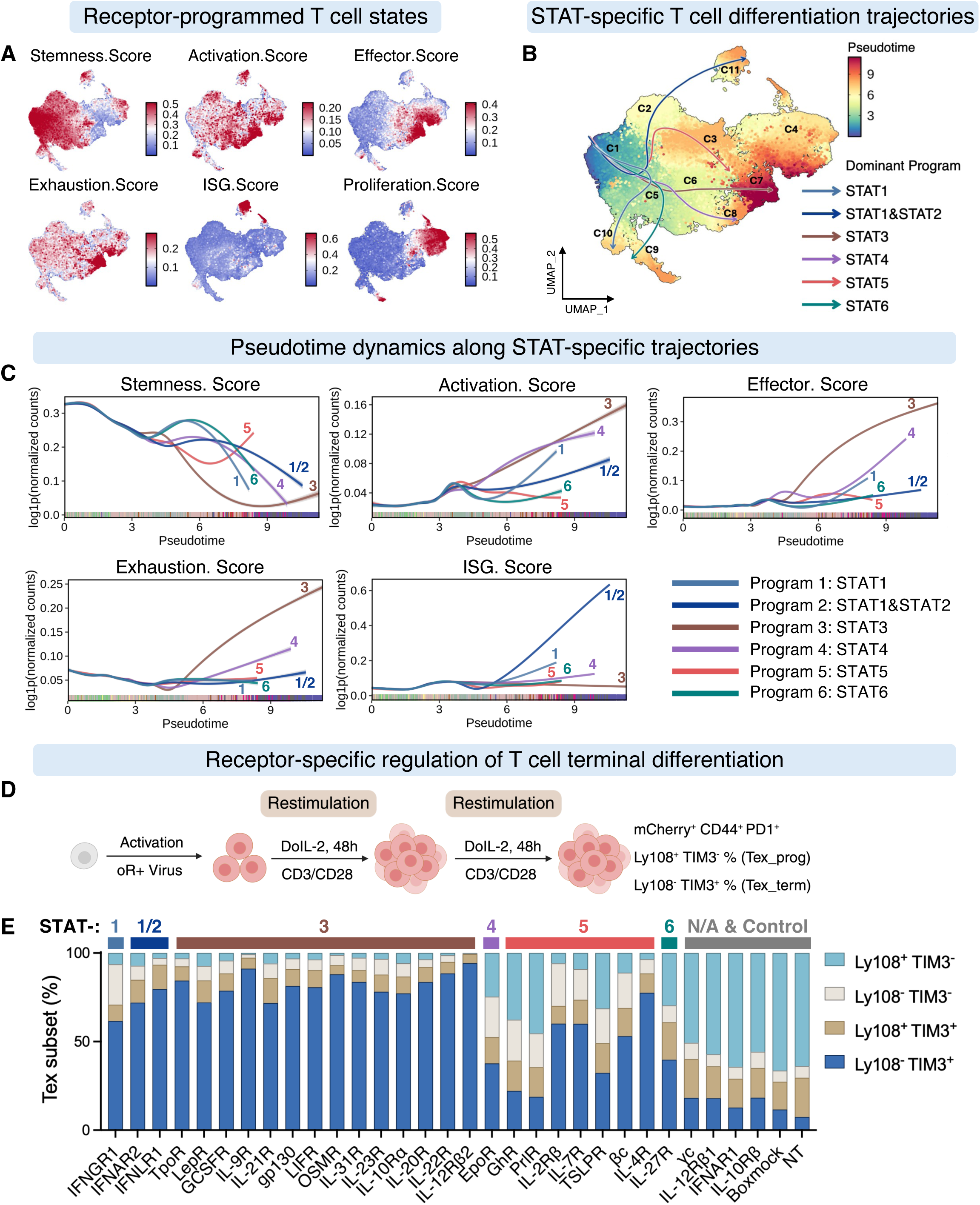
Each STAT signaling program drives T cells along a distinct differentiation trajectory, with STAT3 uniquely programming a functional terminal effector state. (A) Module scores for six functional gene sets, stemness, activation, effector function, exhaustion, interferon-stimulated gene (ISG) response, and proliferation, overlaid on the UMAP, showing which functional programs are enriched in each transcriptional cluster. (B) Slingshot-inferred differentiation trajectories for non-cycling T cells overlaid on the UMAP, colored by pseudotime. Each trajectory reflects a STAT-specific developmental path from early to late states. (C) Dynamics of functional state scores along each inferred differentiation trajectory, plotted against pseudotime. Each panel shows how stemness, effector, exhaustion, and ISG programs evolve as cells progress along a given trajectory. (D) Schematic of the *in vitro* T cell terminal differentiation protocol, in which repeated CD3/CD28 stimulation drives progressive terminal differentiation over multiple rounds of antigen exposure. (E) Frequencies of distinguishing progenitor exhausted T cells (Tex_prog; CD44^+^PD-1^+^Ly108^+^TIM-3^−^) from terminally exhausted T cells (Tex_term; CD44^+^PD-1^+^Ly108^−^TIM-3^+^) populations across all receptor conditions after repeated stimulation, ordered by dominant STAT category. See also **Figure S6** and **S7**.

Although all six trajectories shared a common theme, stemness declining as activation, effector function, and exhaustion rose with pseudotime, the rate and extent of these changes differed dramatically between STAT programs (**Fig. 4C**). STAT3-dominant trajectories began in canonical Tscm states at early pseudotime but underwent the most pronounced transition toward terminal differentiation, ultimately reaching the highest combined activation, effector, and exhaustion scores. In contrast, STAT5-dominant trajectories preserved stemness throughout pseudotime and showed the slowest progression toward terminal differentiation, consistent with the well-established role of STAT5 cytokines like IL-7 and IL-15 in maintaining memory T cell pools. STAT4 drove strong activation but comparatively little effector or exhaustion programming, while STAT6 had minimal engagement with any of these programs. ISG activity was almost entirely restricted to STAT1/STAT2-driven trajectories, and notably, STAT1-only receptors (type II interferon) drove substantially weaker ISG responses than STAT1/STAT2-activating receptors (type I/III interferon), revealing functionally distinct transcriptional programs even within the interferon family (**Fig. 4C**).

Because STAT3-, STAT4-, and STAT5-dominant trajectories all converge toward neighboring regions of the UMAP, we examined the specific genes enriched at each trajectory’s endpoint to understand whether their terminal states are truly distinct (**Fig. S6H**). STAT3-driven terminal cluster (C7) exhibited higher expression of effector molecules (*Prf1*, *Gzma*, *Gzmb*) together with multiple inhibitory molecules (*Havcr2*, *Entpd1*, *Cd244a*, *Cd38*, *Il10*, and *Spp1*). However, the canonical terminal exhaustion-associated transcription factor TOX remained low in C7, suggesting that these cells do not represent a fully dysfunctional exhaustion state **(Fig. S6F**). In contrast, STAT4-driven terminal cells (C8), by contrast, were enriched for activation and inflammatory markers (*Cd28*, *Il18r1*, *Il18rap*, *Gbp2*), while STAT5-driven terminal cells retained a clearly stem-like signature (*Tcf7*, *Ccr7*, *Bach2*, *Slamf6*).

To test whether these computationally inferred trajectories predict functional outcomes, we subjected engineered T cells to repeated rounds of antigen receptor stimulation to model chronic antigen exposure (**Fig. 4D**). The results closely matched the trajectory predictions, STAT3- and STAT4-dominant receptors generated the highest frequencies of terminally exhausted T cells (Tex_term), whereas STAT5-dominant receptors preferentially retained progenitor exhausted cells (Tex_prog) (**Fig. 4E, Fig. S7A**). Strikingly, despite their exhaustion-like phenotype, STAT3-programed T cells remained highly functional, showing enhanced production of TNFα, IFNγ, and granzyme B (**Fig. S7B-C**). Although inhibitory receptors were highly expressed, TOX remained low even after repeated stimulation (**Fig. S7D-E**), and inhibitory receptor expression including TIM-3 could be induced by cytokine receptor signaling alone even in the absence of TCR engagement (**Fig. S7F-G**). These observations indicate that STAT3 programs a functional terminal effector state that exhibits exhaustion-associated features but is mechanistically distinct from true dysfunctional exhaustion. These results demonstrate that quantitative STAT signaling biases are not interchangeable, and that each STAT program instructs a distinct differentiation trajectory that drives T cells into functionally unique terminal states.

### TF regulatory programs across all cytokine receptor driven T cell states

Because STATs are transcription factors that act through broader gene-regulatory networks, we next asked how distinct STAT inputs are translated into the transcription-factor (TF) programs defining each T cell state. As a first step, we validated that computationally inferred STAT activity from our single-cell RNA sequencing data, using CollecTRI-derived TF regulons^48^, accurately reflects the actual signaling measured experimentally by phospho-STAT flow cytometry across all 30 receptor conditions. The two measurements were strongly correlated for every STAT family member (**Fig. 5A**) At the cluster level, inferred STAT activity progressively increased with the degree of differentiation, further supporting STAT signaling as a key driver of T cell fate progression (**Fig. S8A**). Across receptor conditions, STAT1 and STAT2 were selectively activated by interferon receptors, whereas STAT3, STAT4, STAT5, and STAT6 exhibited substantial overlapping activities. Notably, STAT3-dominant receptors displayed the broadest cross-activation across STAT family members, implying extensive coupling within the STAT signaling network (**Fig. S8B**).

**Figure 5.**
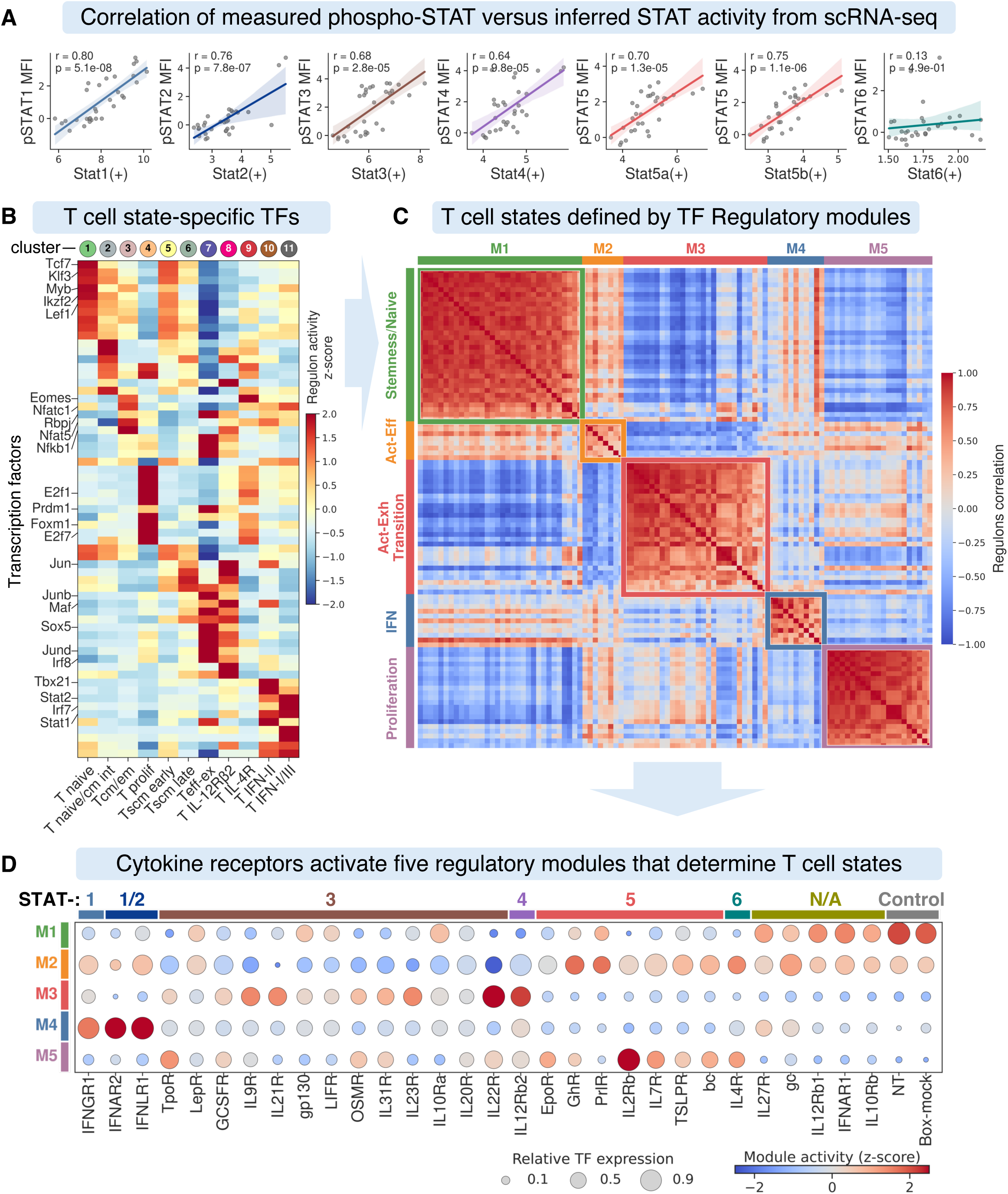
Distinct STAT inputs are decoded into specific transcription factor regulatory circuits, establishing a direct wiring map from receptor signal to gene-regulatory program. (A) Validation of computationally inferred STAT activity against experimentally measured pSTAT levels. Pearson correlation coefficients and p values are shown for each STAT, confirming that inferred activities accurately reflect measured phosphorylation. (B) Heatmap of cluster-specific transcription factor (TF) expression across annotated T cell clusters. Top regulons were selected based on regulon specificity score (RSS). Color indicates scaled TF/regulon activity (z-score). (C) Pairwise regulon correlation heatmap revealing co-regulated TF modules. Regulons are grouped into modules by hierarchical clustering and ordered within each module by mean pairwise correlation. Color indicates Pearson correlation coefficient; colored bars above identify module identity. (D) TF regulatory module activity across all receptor conditions. Each module represents a co-regulated set of TFs; activity is scored as the mean regulon AUC for TFs within each module per receptor condition. Color indicates z-scored module activity; dot size reflects relative TF expression. See also **Figure S8**.

To further delineate the global TF regulatory network, we applied SCENIC^49^ analysis, a computational framework that identifies active gene regulatory networks by linking transcription factor expression to the genes each factor controls (its “regulon”), to infer active regulons across the T cell atlas and ranked cluster-specific TFs based on regulon specificity scores (RSS) (**Fig. S8C**). By integrating regulon activity with TF expression, we identified key TFs defining distinct cluster-specific functional T cell states (**Fig. 5B**). Naïve-like (C1) and early Tscm states (C5) were governed by well-established stemness regulators, Tcf7, Lef1, and Klf3, consistent with their undifferentiated character. As cells differentiated, state-specific TF programs emerged that closely reflected the underlying STAT signaling. The STAT3-driven Teff-ex state (C7) preferentially activated AP-1-associated regulons (Junb and Jund) together with terminal differentiation- and exhaustion-associated regulons (Prdm1, Maf, Sox5, and Irf8), whereas STAT5-associated states displayed stronger NFAT and NF-κB activity together with Eomes. IFN receptor-programmed states were distinguished by selective activation of Stat1/2- and Irf7-centered regulons.

Because transcription factors function cooperatively rather than independently, we next asked whether the individual TF programs identified above organize into higher-order cross-talking regulatory circuits. We computed pairwise correlations between all TF regulon activities across every cell in the atlas, which revealed a highly organized modular architecture with five distinct TF co-activity modules (**Fig. 5C**). These modules correspond to recognizable functional programs: a stemness/naïve module (M1) anchored by Tcf7, Lef1, and Bach2; an activation and effector differentiation module (M2) built around Nfatc1, Nfat5, Nfkb1, and Eomes; an activation-to-exhaustion transition module (M3) dominated by AP-1 TFs alongside Prdm1, Maf, and Irf8; an interferon-response module (M4) defined by Stat1, Stat2, and Irf7; and a proliferation-associated module (M5) marked by cell cycle regulators. These modules showed not only strong internal co-activity but also mutually exclusive patterns between modules (**Fig. S8D-E**), indicating that they represent competing regulatory programs. At the receptor level, TF module activities closely mirrored the functional transcriptional programs identified by gene expression analysis. Receptors with minimal STAT activation retained the stemness/naïve module (M1) as their dominant program. STAT5-dominant receptors most strongly engaged the activation-effector (M2) and proliferation (M5) modules, consistent with their role in driving T cell expansion while preserving functional competence. STAT3- and STAT4-dominant receptors preferentially activated the activation-to-exhaustion transition module (M3), in line with their trajectory toward terminal states. The interferon-response module (M4) was highly restricted, appearing only in the three interferon receptor conditions (**Fig. 5D**).

Finally, to link STAT signaling inputs to TF regulatory outputs, we directly correlated receptor-specific phospho-STAT measurements with TF module activities across all conditions (**Fig. S8F**). This analysis revealed a specific wiring logic: STAT1 and STAT2 were tightly coupled to the interferon-response module (M4); STAT3 and STAT4 were most strongly associated with the activation-to-exhaustion transition module (M3); and STAT5 was the primary driver of the proliferation module (M5). Together, this establishes a quantitative input-output map in which specific STAT combinations are reliably decoded into specific TF regulatory circuits, explaining at the gene-regulatory level how the coarse-grained STAT signaling fingerprints produce the nuanced phenotypically distinct T cell states observed throughout this study.

### Interwoven cytokine receptor signaling balances T cell proliferation and differentiation

Because receptors activate multiple STATs and non-STAT pathways together, we next asked how each receptor’s signaling balance determines two properties central to immunotherapy: 1-how much a T cell expands, and 2-how stem-like it remains. To move beyond the 24-hour transcriptomic snapshot used to build the atlas, we cultured engineered T cells under sustained DoIL-2 stimulation for three days without any T cell receptor (TCR) signaling input, then measured cell proliferation and the frequency of naïve/stem-like memory T cells (CD44⁻CD62L⁺, hereafter naïve/Tscm) for each receptor condition (**Fig. 6A**). This revealed a receptor-dependent hierarchy of proliferative capacity. STAT5-dominant receptors, IL-2Rβ, βc, IL-7R, EpoR, and TSLPR, drove the strongest expansion, consistent with the established role of STAT5 cytokines in supporting T cell survival and proliferation. Receptors engaging a mixed STAT1/3/4/5 program (TpoR, GCSFR, OSMR, IL-31R, and IL-9R) fell into a middle tier, while receptors primarily activating STAT1 and STAT3 (LepR, gp130, IL-10Rα) supported the weakest proliferation (**Fig. 6B**). Strikingly, the pattern for stem-like memory T cells was the reverse: conditions that drove the most proliferation retained the fewest naïve/Tscm-like cells population, and conditions that drove the least proliferation preserved the most (**Fig. 6B-C**). Tscm-associated surface markers Sca-1 and CD95 tracked the same way (**Fig. S9A-B**). This reveals a trade-off by cytokine receptor signaling where proliferative expansion and stem-like memory preservation are competing outcomes, determined by each receptor’s unique STAT signaling mosaic.

**Figure 6.**
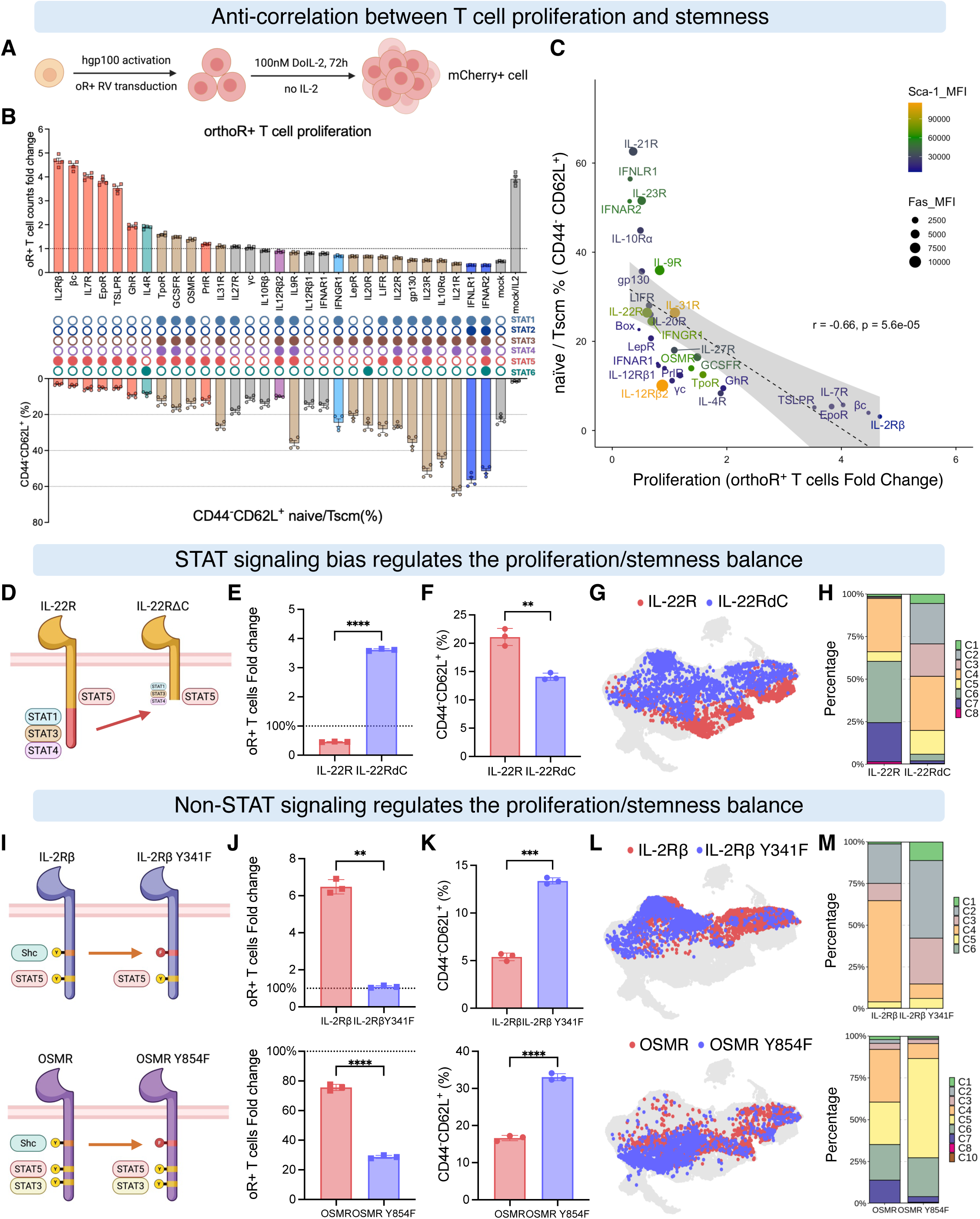
Each cytokine receptor establishes a distinct balance between T cell proliferative expansion and stem-like memory maintenance through the interplay of STAT and non-STAT signaling pathways. (A) Schematic of the *in vitro* proliferation and phenotyping assay. T cells expressing each of 30 orthogonal receptor constructs were stimulated with DoIL-2 (100 nM) for 3 days, then analyzed for cell counts and phenotype. (B) Coupled bar plots linking T cell proliferative expansion and stem-like memory frequency across all receptor conditions. Top: fold change in cell counts. Middle: Major downstream STAT category for each receptor. Bottom: frequency of stem cell-like memory (Tscm; CD44^−^CD62L^+^) T cells. Data shown as mean ± SD (n = 4). (C) Inverse correlation between proliferative expansion and Tscm frequency across all receptor conditions. Each point represents one receptor; Pearson r and p value are shown. Dot color indicates Sca-1 MFI; dot size reflects CD95 (Fas) MFI. (D) Schematic of the IL-22R C-terminal truncation variant (IL-22RdC), which selectively attenuates STAT1/3/4 activity while preserving STAT5 signaling, allowing the STAT1/3/4 contribution to IL-22R function to be dissected. (E-F) Effect of STAT3/4 attenuation on proliferation (E) and Tscm frequency (F) in T cells expressing IL-22R versus IL-22RdC. Both metrics shift toward a more STAT5-like profile upon STAT1/3/4 loss. (G) Transcriptional consequences of STAT1/3/4 attenuation. scRNA-seq data from IL-22R and IL-22RdC T cells are co-embedded in the full T cell UMAP atlas, showing a shift in cluster occupancy. (H) Stacked bar plot of cluster distributions for IL-22R and IL-22RdC, quantifying the transcriptional state shift in (G). (I) Schematic of point-mutant receptor variants that abolish non-STAT signaling without affecting STAT activity: IL-2Rβ(Y341F) removes the Shc-binding site; OSMR(Y854F) does the same for OSMR. These variants isolate the contribution of non-STAT signaling (MAPK, PI3K) to T cell proliferation and stemness. (J-K) Effect of non-STAT signaling ablation on proliferation (J) and Tscm frequency (K) in T cells expressing IL-2Rβ or IL-2Rβ(Y341F), and OSMR or OSMR(Y854F). Non-STAT signaling primarily drives proliferation, with modest effects on stemness. (L) scRNA-seq co-embedding of IL-2Râ, IL-2Râ(Y341F), OSMR, and OSMR(Y854F) T cells in the full atlas UMAP, showing that non-STAT signaling ablation shifts cells toward more stem-like clusters. (M) Stacked bar plots of cluster distributions for these four conditions, quantifying the transcriptional state shifts in (L). See also **Figure S9**.

IL-22R provided a notable exception that let us dissect the contribution of STAT crosstalk to this balance. Despite inducing stronger STAT5 activation than EpoR or IL-7R, IL-22R supported substantially weaker T cell proliferation. We speculated that IL-22R also strongly activates STAT1, STAT3, and STAT4, which appear to antagonize the proliferative output of STAT5. To verify our hypothesis, we analyzed a C-terminal truncated IL-22R (IL-22RdC)^50^, that retains normal STAT5 activation but loses most of its STAT1/3/4 signaling (**Fig. 6D, Fig. S9C**). This single change was sufficient to substantially increase T cell expansion and reduce the proportion of naïve/Tscm-like cells, shifting the balance point toward proliferation (**Fig. 6E-F, Fig. S9D-E**). Single-cell transcriptomic analysis showed that the underlying states change shifted cells away from the differentiated Tscm-late and exhausted-like effector clusters (C6 and C7) and toward STAT5 programed central/effector memory states (C2 and C3) (**Fig. 6G-H**). These results directly demonstrate that receptor-intrinsic STAT signaling mosaics quantitatively tune the balance between T cell proliferation and stemness within a single receptor.

We then asked whether non-STAT pathways contribute similarly by introducing point mutations, IL-2Rβ(Y341F) and OSMR(Y854F), that abolish the Shc docking site and block MAPK/PI3K activation while leaving STAT signaling intact (**Fig. 6I, Fig. S9F**). Removing this non-STAT signaling arm from either receptor substantially reduced T cell expansion and shifted the population toward more stem-like states (**Fig. 6J-K, Fig. S9G-I**), demonstrating that the MAPK/PI3K arm quantitatively cooperates with STAT signaling to drive the proliferative side of the balance. Single-cell transcriptomics confirmed that the proliferating cluster (C4) was sharply diminished in both mutant conditions, while distinct stem-like clusters were enriched, naïve/central memory intermediates (C2) for the IL-2Rβ mutant, and early Tscm cells (C5) for the OSMR mutant, suggesting non-STAT pathways cooperate with STAT signaling to fine-tune the reciprocal balance between T cell proliferation and stemness (**Fig. 6L-M**).

In sum, each receptor’s precise balance of STAT and non-STAT activation orchestrates a distinct reciprocal relationship between T cell proliferative expansion and memory-state differentiation.

### Systematic *in vivo* profiling of T cell antitumor efficacy across all cytokine receptors

Having established these signaling-encoded properties *in vitro*, we next asked whether they shape T cell function *in vivo* in tumor-bearing mice, using an unbiased screen that evaluated all receptor programs in one experiment. Each engineered T cell expressed a single receptor linked to a unique genetic barcode, allowing simultaneous tracking of every receptor-defined population in one host (**Fig. 7A**). As validation, barcode sequencing following 4 days of *in vitro* DoIL-2 stimulation faithfully recapitulated the T cell proliferation hierarchy observed in individual receptor assays, with IL-2Rβ, βc, EpoR, and IL-7R preferentially enriched, whereas IFNAR2, IFNLR1, IL-21R, IL-22R, and the Box control remained depleted (**Fig. S10A-B**).

**Figure 7.**
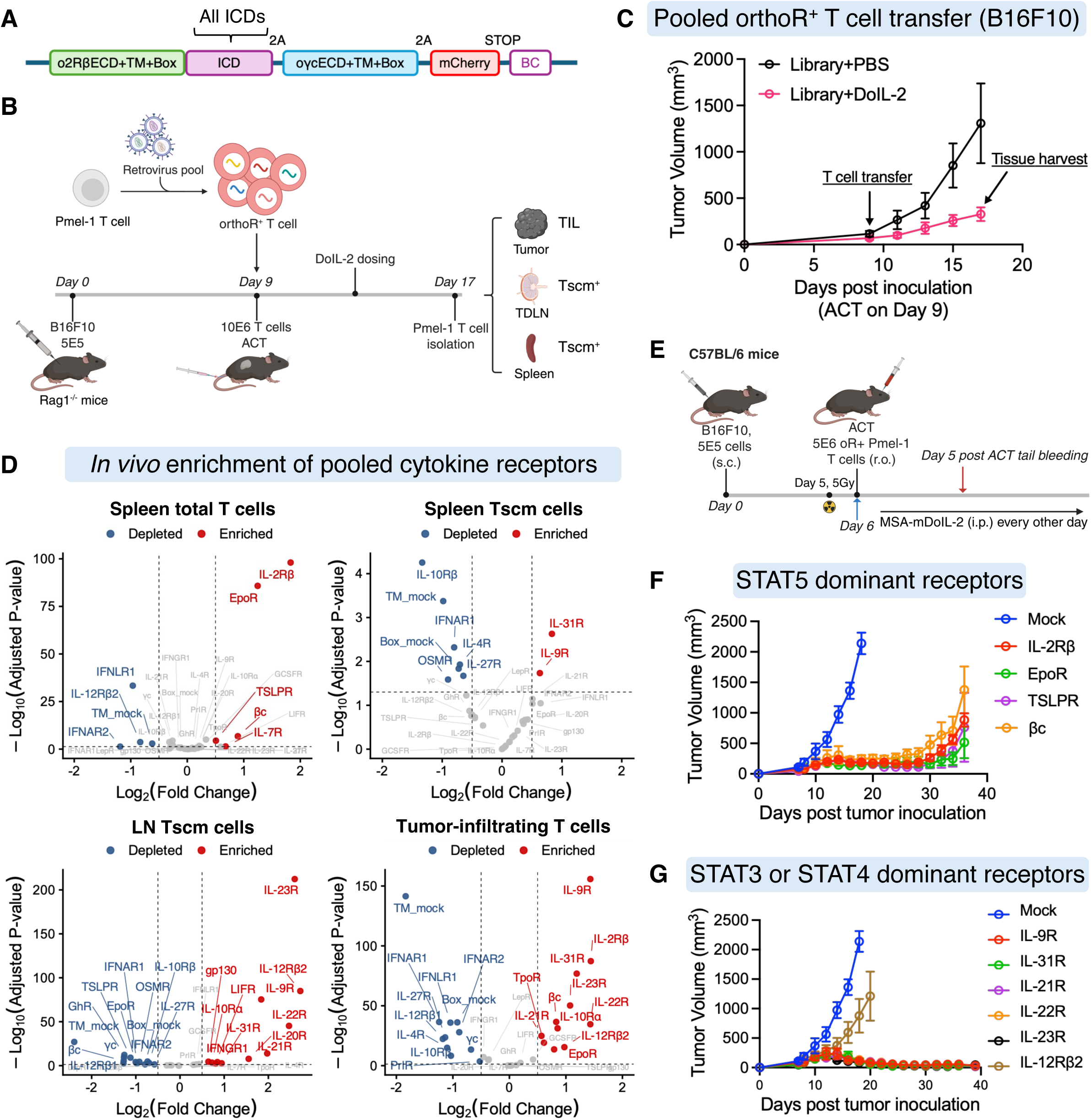
*In vivo* pooled screening identifies STAT3-dominant receptors as conferring durable, curative antitumor efficacy. (A) Schematic of barcoded orthogonal receptor retroviral vectors. Each receptor ICD is paired with a unique DNA barcode, allowing the relative abundance of each receptor condition to be tracked by sequencing in complex pooled experiments. (B) Timeline of *in vivo* pooled selection. Pmel-1 T cells transduced with the barcoded receptor library were adoptively transferred into B16F10 tumor-bearing mice and treated with DoIL-2. Transferred T cells were subsequently isolated from tumor, tumor-draining lymph nodes, and spleen for barcode sequencing to identify which receptor programs were selectively enriched in each compartment. (C) B16F10 tumor growth curves in mice that received the barcoded library transfer and DoIL-2 or control treatment, confirming that the pooled transfer produces expected antitumor activity. Mean ± SD. (D) Volcano plots showing differential barcode enrichment across T cell populations from *in vivo* selection, total T cells in spleen and tumor (TIL), and Tscm populations in spleen and lymph nodes (LN). Receptors enriched in TILs and Tscm compartments identify signaling programs that support tumor infiltration and long-term persistence. (E) Experimental timeline for individual validation of top-ranked receptors enriched in TILs from the pooled screen. Pmel-1 T cells expressing each selected receptor were adoptively transferred into B16F10 tumor-bearing mice and treated with mouse DoIL-2. (F-G) B16F10 tumor growth curves from individual validation experiments for receptor conditions with dominant STAT5 (F) or STAT3, STAT4 (G) signaling. STAT3-dominant receptors consistently achieved durable tumor eradication, while STAT5-dominant receptors failed despite robust expansion. See also **Figure S10**.

We applied the pooled screen in the B16F10 model, transferring Pmel-1 T cells expressing each barcoded receptor into tumor-bearing mice and dosing DoIL-2 (**Fig. 7B**). DoIL-2 dosing significantly improved tumor control compared to PBS, reflecting the collective activity of all receptor conditions (**Fig. 7C**). Isolating total and Tscm T cells (CD44⁻CD62L⁺Sca-1⁺) from spleen and tumor-draining lymph nodes (LN) and total tumor-infiltrating T cells (TIL) (**Fig. 7B**) revealed compartment-specific patterns. Splenic total T cells mirrored *in vitro* proliferation, with STAT5-dominant receptors (IL-2Rβ, EpoR, βc) dominating expansion (**Fig. 7D, Fig. S10C**). The splenic Tscm pool instead favored receptors with mixed STAT1/3/4/5 signaling (IL-31R, IL-9R); in lymph nodes the shift was stronger, favoring more STAT3-or STAT4-dominant receptors (IL-23R, IL-12Rβ2, IL-22R, IL-21R). Within the tumor, enrichment reflected a hybrid pattern, with Tscm-associated receptors (IL-9R, IL-22R, IL-31R, IL-23R) alongside proliferative receptors (IL-2Rβ, βc, EpoR), unlike the receptor preferences observed in either spleen or lymph nodes (**Fig. 7D, Fig. S10C**), suggesting that distinct cytokine receptor programs contribute differently to antitumor immunity.

To determine which receptor programs ultimately confer therapeutic benefit, receptors enriched within TILs were evaluated individually in the fully murine double-orthogonal system (**Fig. 7E**). All receptors improved tumor control relative to the Box control and therapeutic efficacy differed strikingly according to their signaling programs. Despite their superior systemic expansion and strong tumor infiltration, STAT5-dominant receptors (IL-2Rβ, βc, EpoR, TSLPR) were unable to achieve complete tumor regression in any animal; they delayed tumor growth but could not clear it (**Fig. 7F, Fig. S10D-G**). In contrast, receptors engaging combinatorial signaling with dominant STAT3 activation, IL-9R, IL-31R, IL-21R, IL-22R, and IL-23R, achieved robust, durable, and in many cases curative antitumor responses (**Fig. 7G, Fig. S10D-H**).

Together, these findings show that systemic expansion, stem-like memory maintenance, tumor accumulation, and durable antitumor efficacy are distinct functional properties, each instructed by a different subset of cytokine receptor signaling programs. Among all receptor programs tested, dominant STAT3 signaling was the most effective at mediating durable, curative antitumor immunity, marking it as a key instructive signal for engineering T cell therapies.

## Discussion

Given that over 36 cytokine receptors signal through only six STAT transcription factors, it has been a longstanding puzzle whether, and how cytokine receptor signaling is functionally redundant or selective^51–53^. Prior studies have generally concluded that overlapping pathway usage renders cytokine receptors interchangeable^54–58^, but these conclusions rest on bulk measurements that lack the resolution to distinguish quantitative signaling differences between individual receptors. By engineering a fully orthogonal IL-2/IL-2Rβ/γc platform for interference-free, receptor-by-receptor dissection of JAK-STAT signaling across the entire murine repertoire, we show instead that each receptor generates a characteristic STAT fingerprint, defined by relative stoichiometry and signaling strength, and that even subtle differences in this fingerprint instruct profoundly distinct T cell fates, within an evolutionarily fine-tuned architecture that also constrains functional diversity.

A key finding of our work, that both builds on and extends prior studies^4,54,59–61^, is how sensitively cytokine receptor specificity is influenced by quantitative and combinatorial STAT signaling biases, and that minor differences in these biases instruct profoundly different functional outcomes. Across this landscape, STAT family members exhibit strongly correlated and interwoven activation patterns, particularly among STAT1, STAT3, and STAT4. These correlations arise from structural constraints attributable to the high homology of their SH2 domains^62,63^, which enables engagement with similar phosphorylated docking sites^39^, indicating that STAT signaling operates as a coupled network rather than a set of independent modules. This coupling suggests that the signaling landscape accessible to engineered receptors is fundamentally constrained by the intrinsic combinatorial architecture of STAT signaling. Prior studies have suggested that cytokine responses can be tuned by quantitative differences in STAT activation^37,64^, and that signaling specificity emerges from a combination of receptor abundance^65,66^, ligand affinity^67,68^, signaling kinetics^69,70^, and interplay among STAT family members^59,61,71,72^. By systematically profiling all JAK-associated cytokine receptors within a fully insulated signaling framework, our work extends these concepts and demonstrates that quantitative and combinatorial STAT biases constitute a general organizing principle underlying cytokine receptor specificity.

Single-cell profiling revealed the cellular programs tied to each receptor’s signaling identity: although receptor-level embeddings appear promiscuous, the transcriptional landscape is organized along a STAT-guided architecture. Distinct clusters formed by interferon receptors (IFNAR2, IFNLR1, and IFNGR1) highlight the critical role of STAT2 in differentiating type I/III from type II interferon responses through coordination with STAT1^73,74^. In addition, IL-12Rβ2 also formed a discrete cluster, which may be explained by the fact that although it activates multiple STATs, its transcriptional program is dominated by STAT4 signaling^75^. In contrast, IL-4R, which selectively activates STAT6, occupied a discrete cluster and exhibited a distinct transcriptional program but distinct from the canonical Tc2 phenotype, without GATA3 activation and IL-4/IL-5/IL-13 expression, likely reflecting the absence of TCR stimulation required for Th2/Tc2 lineage specification^76,77^. All remaining receptors organized into a continuous differentiation hierarchy governed by their STAT combinatoric balance. Notably, several shared receptor chains (γc, IFNAR1, IL-12Rβ1, and IL-10Rβ), which possess short intracellular domains and lack well-defined STAT docking motifs, still elicited modest transcriptional responses compared with the mock control, suggesting potential noncanonical or context-dependent signaling^78,79^. Together, these transcriptomic data show that cytokine receptor signaling is not redundant. Each receptor encodes a non-interchangeable T cell fate through a unique hierarchy of dominant and subordinate STAT signaling inputs, and the prior perception of redundancy arose from bulk measurements that lacked the resolution to distinguish these quantitative combinatorial differences.

A notable functional insight revealed in this study is the trade-off between T cell proliferation and stem-like memory maintenance, two critical parameters for adoptive T cell therapy. Across receptor conditions, we observed a strong inverse relationship between proliferative capacity and the generation of Tscm-like populations. This balance is governed by the interplay between STAT5-dominant signaling, which promotes metabolic activity and cell cycle progression, and STAT1/3/4-associated programs, together with non-STAT pathways such as Shc-MAPK signaling, which influence differentiation and memory formation. These findings align with prior strategies that enhance Tscm generation through transient modulation of STAT5^80^, MAPK^81^, PI3K-AKT^82,83^, or mTOR^84^ signaling, as well as activation of Wnt/β-catenin^85^ pathways, and provide a unified receptor-level explanation for these observations.

*In vivo* pooled screening and individual therapeutic testing revealed that systemic expansion, Tscm maintenance, tumor accumulation, and curative efficacy are distinct properties prioritized by different receptor programs. Despite driving robust *in vivo* expansion and tumor infiltration, STAT5-biased receptors failed to generate durable tumor control. In contrast, receptors enriched for STAT3-dominant combinatorial signaling (including IL-9R, IL-21R, IL-22R, IL-23R and IL-31R) exhibited superior curative efficacy. Mechanistically, STAT3 signaling defined a continuous differentiation program spanning from stem-like Tscm cells to a distinct terminal effector population exhibiting partial exhaustion-associated features. Expression of Tox, a canonical TF of dysfunctional exhaustion, was reduced and negatively correlated with STAT3 signaling. These findings indicate that STAT3-driven exhaustion-like features do not necessarily represent dysfunction, but instead reflect a distinct, cytokine-programmed terminal differentiation state with preserved cytotoxic capacity. This observation is consistent with emerging evidence that cytokines such as IL-10, IL-21, IL-27, and IL-4 can reinvigorate exhausted T cell states within tumors^86–90^, whereas our results further suggest that STAT3 signaling intrinsically facilitates the generation of terminally differentiated effector populations with exhaustion-associated features. We propose that this differentiation heterogeneity is functionally advantageous: Tscm-like cells sustain long-term persistence, while terminally differentiated STAT3-dependent effectors provide immediate cytotoxic activity, together enabling durable tumor control.

In summary, cytokine receptor signaling selectivity is encoded in fine-tuned quantitative STAT combinatoric fingerprints distributed across a structurally constrained receptor landscape. These findings provide a complete receptor-to-function decoding of cytokine signaling logic and establish principled design rules for engineering cytokine receptors to optimize T cell-based immunotherapies.

### Limitations of the study

First, although we profiled single-cell transcriptomes of mouse T cells in response to signaling from all natural cytokine receptors under *in vitro* conditions, the landscape of T cell states shaped by concurrent TCR stimulation remains uncharacterized, particularly in tumor-infiltrating T cells undergoing chronic antigen exposure. Second, while STAT-associated transcriptional signatures were resolved at the single-cell level, the direct gene regulatory programs governed by individual STAT factors remain undefined. Integrating chromatin accessibility approaches, such as ATAC-seq, would enable reconstruction of STAT-centric cis-regulatory networks with higher resolution. Finally, whether receptor-specific signaling profiles and transcriptional programs are conserved between murine and human systems remains to be determined, which will be critical for supporting translational applications.

## Acknowledgments

This work was supported by NIH-RO1-AI51321 (K.C.G.), Parker Institute for Cancer Immunotherapy (K.C.G.), Stanford-UCSF Weill Cancer Research Fund, Ludwig Institute, K.C.G. is an Investigator of the Howard Hughes Medical Institute. We acknowledge the Animal care facilities at Stanford, Stanford Shared FACS Facility for their technical assistance. Schematics and diagrams were created using BioRender.

## Author contributions

K.C.G. conceived the study, supervised the experiments, and collaborated with the other authors on the manuscript. P.T. designed and conducted the experiments, analyzed the data, and wrote the manuscript. R.R. and A.K contributed to the analysis of scRNA-seq data. H.J. and Y.Z. assisted with *in vivo* mouse experiments. L.S. supported the design and construction of orthogonal receptor plasmids. K.J. assisted with IL-2R complex structure analysis and mutant library design. All authors contributed to manuscript preparation.

## Data availability

The raw and processed scRNA-seq data generated in this study will be deposited for public availability in the GEO database upon publication. The code used for RNA-seq data analysis will also be shared publicly at that time. Any additional information required to reanalyze the data reported in this paper is available from the lead contact upon request.

## Methods

### Experimental model and study participant details Mice

Five-to eight-week-old Thy1.2^+^ C57BL/6 (C57BL/6J), Thy1.1^+^ C57BL/6 (B6.PL-Thy1a/CyJ), Thy1.1^+^ Pmel-1 TCR-transgenic mice (B6.Cg-Thy1a/Cy Tg(TcraTcrb)8Rest/J) and Rag1 KO mice (B6.129S7-Rag1tm1Mom/J) were originally purchased from the Jackson Laboratory and maintained in the Stanford University-Lorry Lokey (SIM1) Facility. All animal experiments were conducted in accordance with the guidelines of the protocol (ID: 32279) approved by the Institutional Animal Care and Use Committee (IACUC), Administrative Panel on Laboratory Animal Care (APLAC) at the Stanford University.

### Cell lines and cell culture

The B16-F10 mouse melanoma cell line and 293T cell was purchased from ATCC and cultured with complete DMEM medium (high-glucose DMEM supplemented with 10% FBS, 2 mM GlutaMAX, 10 mM HEPES, and penicillin-streptomycin). Platinum-E (Plat-E) and Platinum-GP (Plat-GP) retroviral packaging cell lines were purchased from Cell Biolabs and cultured with complete DMEM medium. STAT5 luciferase reporter YT cell line was derived from YT-1 cell line by stable transduction with pGL4.52 vector (Promega) and cultured with complete RPMI 1640 medium (RPMI 1640 supplemented with 10% FBS, 50 μM 2-mercaptoethanol, non-essential amino acids, sodium pyruvate, 2 mM GlutaMAX, 10 mM HEPES, and penicillin-streptomycin). Expi293F cells were purchased from Gibco and cultured in suspension with Expi293 Expression Medium. All related cell culture reagents were purchased from Gibco.

### Primary T cells preparation and culture

Primary mouse total T cells or CD8^+^ T cells were isolated from spleen and lymph nodes of six-to eight-week-old mice using Pan T Cell Isolation Kit II (Miltenyi) or CD8a^+^ T Cell Isolation Kit (Miltenyi) and cultured in complete RPMI 1640 medium supplemented with 100 U/mL mouse IL-2 (Miltenyi). Human PBMC were isolated from LRS chambers (Stanford Blood Center) and cryopreserved until time of use. Human T cells were isolated from thawed PBMC using EasySep™ Human T Cell Isolation Kit (STEMCELL) and cultured in complete RPMI 1640 medium supplemented with 100 U/mL human IL-2 (PeproTech).

### Yeast strains and culture

Saccharomyces cerevisiae yeast strain (EBY100, ATCC) was cultured at 30 °C in YPD media (Sigma) and selected in synthetic defined media containing glucose and casamino acids (SDCAA): 1L deionize water with 20 g dextrose/glucose (Sigma), 6.7 g yeast nitrogen base (RPI), 5 g Bacto casamino acids (Gibco), 10.4 g sodium citrate (Sigma), and 6.4 g citric acid monohydrate (Sigma), pH 4.5. Surface displayed protein was induced at 20 °C in galactose containing media (SGCAA) which resembles SDCAA but includes 20 g galactose (Sigma) instead of dextrose/glucose.

### Method details Protein production

DNA encoding human IL-2 and its variants, mouse IL-2 and its variants, human IL-2Rβ extracellular domain (ECD), and human IL-2-IL-2Rβ ECD fusion protein were cloned into the mammalian expression vector pD649, which includes a C-terminal Avi-tag for biotinylation and 6x His tag for affinity purification. DNA encoding mouse serum albumin (MSA) was cloned into the N-terminal of the pD649 constructs described above to express MSA fusion protein. Mammalian expression DNA constructs were transfected into Expi293F cells using the Expi293 Expression System (Gibco) for secretion and purified from the clarified supernatant by nickel affinity resin (Ni-IMAC, Thermo Fisher Scientific) followed by size-exclusion chromatography with a Superdex-200 column (Cytiva) and formulated in sterile phosphate-buffered saline (PBS). Endotoxin was removed using the Proteus NoEndo HC Spin column kit (VivaProducts) and endotoxin removal was confirmed using the Pierce LAL Chromogenic Endotoxin Quantification Kit (Thermo Fisher Scientific). Purified proteins with Avi-tag were biotinylated *in vitro* with BirA ligase (produced in house) and then repurified from the reaction mixture by size exclusion chromatography. Proteins were concentrated and stored at -80 °C until use.

### Yeast display of γc extracellular domain and binding validation

Human γc extracellular domain was displayed on the surface of the yeast EBY100 by fusion to the N-terminal of Aga2 using the pYAL vector harboring a HA epitope tag for displayed protein detection. Briefly, competent EBY100 was electroporated with plasmid encoding yeast-displayed hγc-ECD and recovered overnight in synthetic defined medium with casamino acids (SDCAA) selection medium at 30 °C. Transformed yeast were passaged once in SDCAA, and yeast cultures in log phase were pelleted and resuspended at an OD600 (optical density at 600 nm) of 1.0 in SGCAA (selective galactose CAS amino acid induction media) induction medium containing 10% SDCAA and cultured for 24-48 hours at 20 °C. Surface expression of functional hγc-ECD was confirmed by FACS by staining yeast with an anti-HA monoclonal antibody (1:100 dilution; Cell Signaling Technology) and a hγc-ECD structure specific nanobody (VHH3) we described before^91^. IL-2/IL-2Rβ/γc ternary complex formation was validated by FACS by staining yeast with pre-mixed human IL-2/IL-2Rβ binary complex (500 nM + 500 nM) and human IL-2-IL-2Rβ fusion protein (500 nM).

### Yeast library generation and selection

The site-directed library of human γc was created by assembly PCR using primers with five degenerate codons (NNK) at the position of Q127/N128/P207/L208/S211. The mutated γc gene PCR product was assembled using PrimeSTAR Max DNA Polymerase (Takara) and an equal molar mixture of designed primer (IDT). The resulting assembled PCR product was gel-purified and electroporated with linearized pYAL vector into EBY-100 yeast to yield a library of about 1 × 10^8^ transformants as described^92^. Selection of yeast clones that specifically bind the mutated IL-2-IL-2Rβ fusion protein was performed using three rounds of magnetic-activated cell sorting (MACS) followed by one round of fluorescence assisted cell sorting (FACS). The first round of selection was performed with 1 × 10^9^ yeast cells and 500 nM biotinylated fusion protein captured on streptavidin microbeads. The final round of positive sorting was performed with 1 × 10^7^ yeast cells stained with 100 nM fusion protein followed by 100 nM streptavidin-AF647 (Invitrogen) to detect binding of biotinylated monomer. Plasmids from enriched yeast clones were isolated using the Zymoprep Yeast Plasmid Miniprep II Kit (Zymo Research) and sequenced for the mutation.

### Construction of orthogonal cytokine receptor expression vectors

cDNA encoding orthogonal IL-2Rβ and engineered orthogonal γc were separately or continually cloned into the lentiviral vector pCDH (System Biosciences) or the retroviral pMSCV vectors containing different fluorescent protein expression (TagBFP, ZsGreen or mCherry) by PCR and isothermal assembly (ITA). Bicistronic expression of both orthogonal receptors at equal level was achieved by a P2A self-cleaving peptide. To construct the orthogonal chimeric cytokine receptors, the transmembrane domain and Box region of all natural cytokine receptors were defined by UniProt and related references^31,32^. The amino acid sequences of all tested orthogonal chimeric cytokine receptors with original or fixed Box regions were listed in the Supplementary Information.

### Retrovirus and Lentivirus production

To generate retrovirus for the transduction of mouse T cells, Plat-E cells (Cell Biolabs) were seeded at 8 × 10^5^ cells per well with 3 mL medium in a 6-well tissue culture treated plate. Following an overnight incubation, fresh media was replenished prior to transfection. For each well of transfection, 5 µg of plasmid (3 µg of pMSCV retroviral expression vector plus 2 µg of pCL-Eco packaging vector) was added to 300 µl of Opti-MEM I Reduced Serum Medium (Gibco), followed by 15 µl of FuGENE® HD transfection reagent (Promega), with gentle vortexing. After a 15-minute incubation, the transfection mixture was gently added to Plat-E cells. Retrovirus supernatant was collected at 48 hours post-transfection and filtered through a 0.45-µm syringe filter (Nalgene). If not used immediately, the virus was frozen for storage at -80°C. To generate retrovirus for the transduction of human T cells, Plat-GP cells (Cell Biolabs) were prepared under the same condition, but virus production was using a different plasmid transfection recipe (3 µg of pMSCV retroviral expression vector plus 2 µg of pLTR-RD114A envelope plasmid).

Lentivirus was produced in HEK293T cells using third-generation packaging vectors. Briefly, HEK293T cells were seeded at a density of 5 × 10^5^ cells per well in a 6-well tissue culture treated plate and allowed to adhere with overnight incubation. Supernatant was removed and replenished with 3 mL low-FBS (5%) DMEM and cells were transfected with 5 µg plasmid at a 4:3:1 ratio of pCDH:psPAX2:pMD2G using FuGENE® HD transfection reagent (Promega) and cultured for 48 hours. Lentivirus supernatant was collected, filtered and stored as above.

### Activation and transduction of primary T cells

Isolated T cells from C57BL/6 mouse were activated with plate-bound anti-mouse CD3ε (5 μg/mL, clone 145-2C11, Biolegend) and soluble anti-mouse CD28 (5 μg/mL, clone 37.51, Biolegend) in complete RPMI 1640 medium supplemented with 100 U/mL mouse IL-2 (Miltenyi) for 24 h. Isolated CD8^+^ T cells from Pmel-1 mouse were activated with 1 µM human gp100 peptide (GenScript) in complete RPMI 1640 medium supplemented with 100 U/mL mouse IL-2 (Miltenyi) for 24 h. One day before transduction, 12-well tissue culture plates were coated with Retronectin (25 μg/mL, Takara) and placed in a 4 °C refrigerator overnight. The following day, plates were washed twice with complete RPMI 1640 medium. Viral supernatant (3 mL) was added to each well and spun at 2500 g for 2 hours at 32°C. After spinning, the virus particles were captured on the plate. Activated T cells (1 × 10^6^ in 2 mL medium) were added to each well after aspirating the viral supernatant and spun at 1000 g for 10 minutes at 32°C. After incubating overnight, T cells were collected and expanded in a fresh medium until further analysis. On day 3, transduction efficiency was assessed based on the expression of fluorescent protein using flow cytometry. Untransduced T cells activated and cultured in parallel were used as control. Mouse T cells were collected for *in vitro* assays or *in vivo* injection three days after spinfection.

Isolated human T cell cells were activated with plate-bound anti-human CD3ε (5 μg/mL, clone OKT-3, Biolegend) and soluble anti-human CD28 (5 μg/mL, clone CD28.2, Biolegend) for 48 h. Activated T cells were collected and transduced with viral supernatant on 12-well plates coated with Retronectin as described above. To co-transduce T cells with two different viruses, a secondary transduction was performed 12 hours later after the first spinfection. Transfected human T cells were collected and expanded in a fresh medium until they returned to a resting state for functional assays.

### Phosphoflow signaling assays of primary T cells

Actively growing mouse or human primary T cells were washed and rested in T cell medium without IL-2 for 12-18 hours before signaling assays. Cells were plated in a 96-well round bottom plate in complete RPMI 1640 medium without FBS prior to the assay. To detect phosphorylation of STAT proteins, Ribosomal Protein S6 (RPS6), cAMP-Response Element Binding Protein (CREB) and mTOR, T cells were stimulated by addition of orthogonal cytokines for 30 minutes at 37°C. To detect phosphorylation of Erk, p38 MAPK, Akt and p90RSK, T cells were stimulated by addition of orthogonal cytokines for 5 minutes at 37°C. The reaction was terminated by fixation with 2.1% paraformaldehyde (BD Cytofix) for 30 minutes at 37°C. Fixed cells were washed and permeabilized with ice-cold methanol (BD Phosflow Perm Buffer III) for 30 minutes on ice or stored at -80°C for later analysis. Cells were washed with staining buffer (PBS containing 2% FBS) before staining with phosphoflow antibodies (BD or CST) for 30 min to 1 hour at 4°C in the dark. Cells were washed and analyzed on a CytoFlex (Beckman Coulter) or Novocyte Quanteon (Agilent Technologies). For the specific pSTAT2 detection in mouse T cells, ectopically transduced human STAT2 was used as a surrogate, and analyzed under the positive gate. Data represent the median fluorescence intensity (MFI), and points were fit to the [agonist] versus dose–response (three parameters) model using Prism 10 (GraphPad).

### Single-cell RNA sequencing

CD8^+^ T cells isolated from Pmel-1 mice were activated and transduced with each orthogonal cytokine receptor individually. Engineered T cells (mCherry^+^) were sorted at 48 hours post transfection. After an overnight resting, T cells were stimulated with 100 nM double ortho-IL-2 for 24 hours to initiate the transcription programs. After the stimulation, live T cells were purified with the Dead Cell Removal Kit (Miltenyi) and fixed using the High-Throughput Evercode Cell Fixation v3 Kit (Parse Biosciences). Fixed cells underwent three rounds of combinatorial barcoding according to the protocol of Evercode Cell WT v3 Kit (Parse Biosciences) and resulted in 8 sublibraries for sequencing. The generated 3′ Gene Expression libraries were sequenced using the NovaSeq X Plus (illumina) with a sequencing depth of > 50K paired-end reads per cell.

### Analysis of single cell RNA-sequencing data

The raw FASTQ files from sub-libraries were demultiplexed into all individual samples using the pipeline from Trailmaker pipeline (Parse Biosciences). Reads were aligned to the mouse reference genome (GRCm39), and gene annotations were based on Ensembl release 109. Gene-level count matrices were generated using default parameters. Initial quality control, including filtering of low-quality cells based on UMI counts, mitochondrial gene content, and doublet detection, was performed using the built-in automated pipeline in TrailMaker (Parse Biosciences).

Raw count matrices were exported for downstream analyses, and low-frequency genes detected in fewer than 0.1% of cells were removed. The top 5,000 highly variable genes (HVGs) were identified using the Scanpy highly variable genes function with the Seurat v3 method and used for subsequent analyses. A latent representation of the data was learned using scVI (scvi-tools v1.4.0). The learned latent space was used for neighborhood graph construction, and cells were clustered using the Leiden algorithm at resolution 0.6. Low-dimensional visualization was performed using Uniform Manifold Approximation and Projection (UMAP) based on the scVI latent representation. Downstream analyses were performed using Python (JupyterLab, Scanpy) and R (Rstudio, Seurat v5), and data visualization was generated using both platforms unless otherwise specified.

Trajectory inference was performed using the Slingshot package in R. Leiden-defined T cell clusters and the scVI-derived low-dimensional embedding were used as inputs to infer lineage trajectories. To minimize cell cycle gene-associated effects on lineage reconstruction, trajectory inference was restricted to non-cycling T cells. The naïve-like T cell cluster C1 was defined as the root population, and lineage trajectories were inferred using Slingshot with default parameters unless otherwise indicated. To quantify major T cell functional programs, gene signature scores were calculated using the UCell R package, which performs rank-based single-cell gene signature scoring, with curated gene sets representing distinct T cell functional states (Supplementary Tables). Dynamic changes in the average expression of genes defining the major T cell functional states were analyzed along Slingshot-derived pseudotime to characterize state transitions during T cell differentiation.

For transcriptional regulatory network analysis, regulon activity was inferred using pySCENIC (v0.12.1). Briefly, gene co-expression networks were constructed using GRNBoost2, and candidate regulons were identified with RcisTarget based on motif enrichment analysis. RcisTarget motif rankings based on the mm10 reference databases (500 bp upstream and 10 kb around TSS) were used. Regulon activity in individual cells was then quantified using AUCell, generating area under the curve (AUC) scores at the single-cell level. Regulon activity scores were used to define transcription factors-driven cell states, and regulon specificity scores (RSS) were calculated to identify cluster-specific transcription factors. Scaled regulon activity was used for visualization. For regulon module analysis, pairwise correlations of regulon AUC scores were calculated across single cells to assess co-activity relationships among transcription factor regulons. Representative regulons were grouped into functional modules based on regulon identity, co-activity patterns, and known biological functions. Regulons within each module were ordered by their mean intra-module correlation, and the resulting regulon-regulon correlation matrix was visualized as a heatmap with module annotations and boundaries. In parallel, STAT activity was inferred from gene expression profiles using a univariate linear model (ULM) based on the Collection of Transcriptional Regulatory Interactions (CollecTRI), which integrates signed TF-target gene interactions from multiple resources. The ULM was applied to quantify STAT activity at the single-cell level, and the resulting scores were used for downstream analysis and visualization.

### Mouse T cell *in vitro* proliferation and immunophenotyping

Actively growing transduced mouse T cells were washed and re-suspended in mouse T cell medium lacking IL-2 and seeded at a density of 50,000 T cells per well (in 100 µL) in a 96-well round bottom tissue culture plate. T cell proliferation was stimulated by addition of 100 µL double ortho IL-2 (200 nM) to a total volume of 200 µL. Before the culture, 100 µL T cells were transferred to a new plate for cell counting as day 0. The remaining 100 µL T cells were cultured for 3 days at 37 °C. On day 3, T cells were collected and counted by FACS using the CytoFLEX equipped with a high throughput sampler as compared with day 0. The total number of live orthogonal cytokine receptor expressing live T cells were gated by staining of Fixable Viability Dye eFluor™ 780 (eBioscience) and expression of mCherry. For the immunophenotyping of T cell stemness, CD44, CD62L, Sca-1 and CD95 antibodies were included in the staining buffer and gated during cell counting.

### Mouse T cell *in vitro* re-stimulation and immunophenotyping

Actively growing transduced mouse T cells were washed and re-suspended in mouse T cell medium lacking IL-2 and seeded at a density of 50,000 T cells per well (in 50 µL) in a 96-well round bottom tissue culture plate. T cells were firstly stimulated by addition of 50 µL medium containing double ortho IL-2 (200 nM) and human gp100 (200nM) to a total volume of 100 µL. On day 2, cells were fed fresh cytokine and peptide in an additional 100 µL volume and cultured for another 2 days as secondary stimulation. After the two rounds of stimulation, T cells were collected for phenotyping by flow cytometry. The total number of live orthogonal cytokine receptor expressing live T cells were gated by staining of Fixable Viability Dye eFluor™ 780 (eBioscience) and expression of mCherry. For the immunophenotyping of terminally differentiated T cells, CD44, PD-1, SLAMF6 (Ly108), TIM-3, LAG-3, CD39, 2B4 and TOX were stained for FACS analysis. To evaluate the cytokine secretion of these restimulated T cells, Cell Stimulation Cocktail plus Protein Transport Inhibitor (eBioscience) was added in the medium for 4-6 hours incubation at 37 °C. Intracellular cytokines (IFNγ and TNFα) staining was subsequently processed using the Cell Fixation/Permeabilization Kit (BD Biosciences).

### Barcoded T cell *in vivo* profiling

Each of the 30 natural cytokine receptors and the mock control cloned in the double orthogonal construct (pMSCV-mCherry) were labeled with a unique 8-bp DNA barcode in the vector backbone prior to 3’-LTR. Each construct was midi-prepped separately and pooled at equal molar ratio. The virus pool was prepared using the same packaging recipe as the individual vector. For the pooled library experiments, activated mouse T cells were infected with 1% v/v virus (MOI = 0.5) to reach a transduction rate of under 15%, ensuring that each cell expressed a single receptor type. Six to eight-week-old Rag1^-/-^ mice were subcutaneously injected with 5 × 10^5^ B16F10 tumor cells in 100 μL of PBS into the right flank. 1× 10^7^ transduced Pmel-1 T cells were sorted and transferred into tumor bearing mice via retroorbital injection on day 9. PBS or MSA-DoIL-2 (15 μg) was administered the same day and every other day for a total of five dosings. Tumor size (length and width) was measured with calipers at each treatment point and volume was calculated as (length × width^2^)/2. On Day 17, transferred T cells from spleen, tumor draining lymph nodes and tumor were isolated and sorted as indicated for the following receptor barcodes analysis. Genomic DNA from 5000 to 50,000 sorted T cells and unselected T cells from the infusion product was extracted using Quick Extract (Lucigen) and used as the template for a two-step PCR strategy. In the first PCR reaction, receptor barcodes integrated in the genome were amplified with the primers designed based on the pMSCV backbone. The DNA product was used as a template for a second PCR reaction to add Illumina-compatible adapters for deep sequencing. The resulting amplicons were sequenced on the Illumina MiSeq system using MiSeq Reagent Kit v3 (150-cycle). DESeq2 was used to normalize reads and calculate fold changes and p-values for paired samples collected before and after *in vivo* selection.

### *In vivo* mouse tumor model

Six to eight-week-old C57BL/6J mice were subcutaneously injected with 5 × 10^5^ B16F10 tumor cells in 100 μL of PBS into the right flank. Mice bearing established tumors were sublethally lymphodepleted by whole body irradiation (5 Gy) on day 5. On day 6, mice were randomized based on average tumor size and received adoptive transfer of 5 × 10^6^ Pmel-1 T cells (50% transduced). T cells were resuspended in 50 μl of PBS per mouse and administered by retroorbital intravenous injection. MSA-DoIL-2 (15 μg) in 100μl of PBS was administered the same day and every other day until day 20. Peripheral blood (10 μL) was collected using Microhematocrit Capillary Tubes (Fisherbrand) at indicated time points from the tail vein for quantification of adoptively transferred Pmel-1 T cells by flow cytometry. Tumor size (length and width) was measured with calipers three times a week and volume was calculated as (length × width^2^)/2. Post-therapy survival of mice was monitored for at least 60 days post-tumor inoculation. Mice were euthanized when the total tumor volume exceeded 2,000 mm^3^ or reached the morbidity criteria, as per APLAC guidelines.

### Statistical analysis

Statistical analysis was performed using GraphPad Prism v10, except where indicated. All values and error bars are shown as mean ± SEM. Comparisons of two groups were performed by using two-tailed unpaired Student’s t test. Comparisons of multiple groups were performed by using one-way analysis of variance (ANOVA) with Tukey’s multiple-comparisons test unless otherwise indicated. Experiments that involved repeated measures over a time course, such as tumor growth were performed by using two-way ANOVA with Tukey’s multiple-comparisons post-test. Survival data were analyzed using the Log-rank (Mantel-Cox) test. P-values were considered significant if less than 0.05, indicated as ∗ p <0.05, ∗∗ p <0.01, ∗∗∗ p <0.001, and ∗∗∗∗ p <0.0001. No statistically significant (NS) differences were considered when P-values were larger than 0.05.

## Supplemental information

Table S1. Amino acid sequences of all orthogonal cytokine receptors.

Table S2. Cluster-specific differential gene expression analyses of integrated scRNA-seq data.

Table S3. Gene sets used to identify T cell functional signatures.

Table S4. Cluster-specific transcription factor regulons identified by SCENIC and ranked by regulon specificity scores (RSS).

## Declaration of generative AI and AI-assisted technologies in the writing process

During the preparation of this work, the authors used Claude and ChatGPT to improve the readability of the manuscript and to assist with the analysis and visualization of single-cell RNA sequencing data. The authors reviewed and edited the output as needed and take full responsibility for the content of the published article.

## Supplemental Figures and Legends

**Figure S1.**
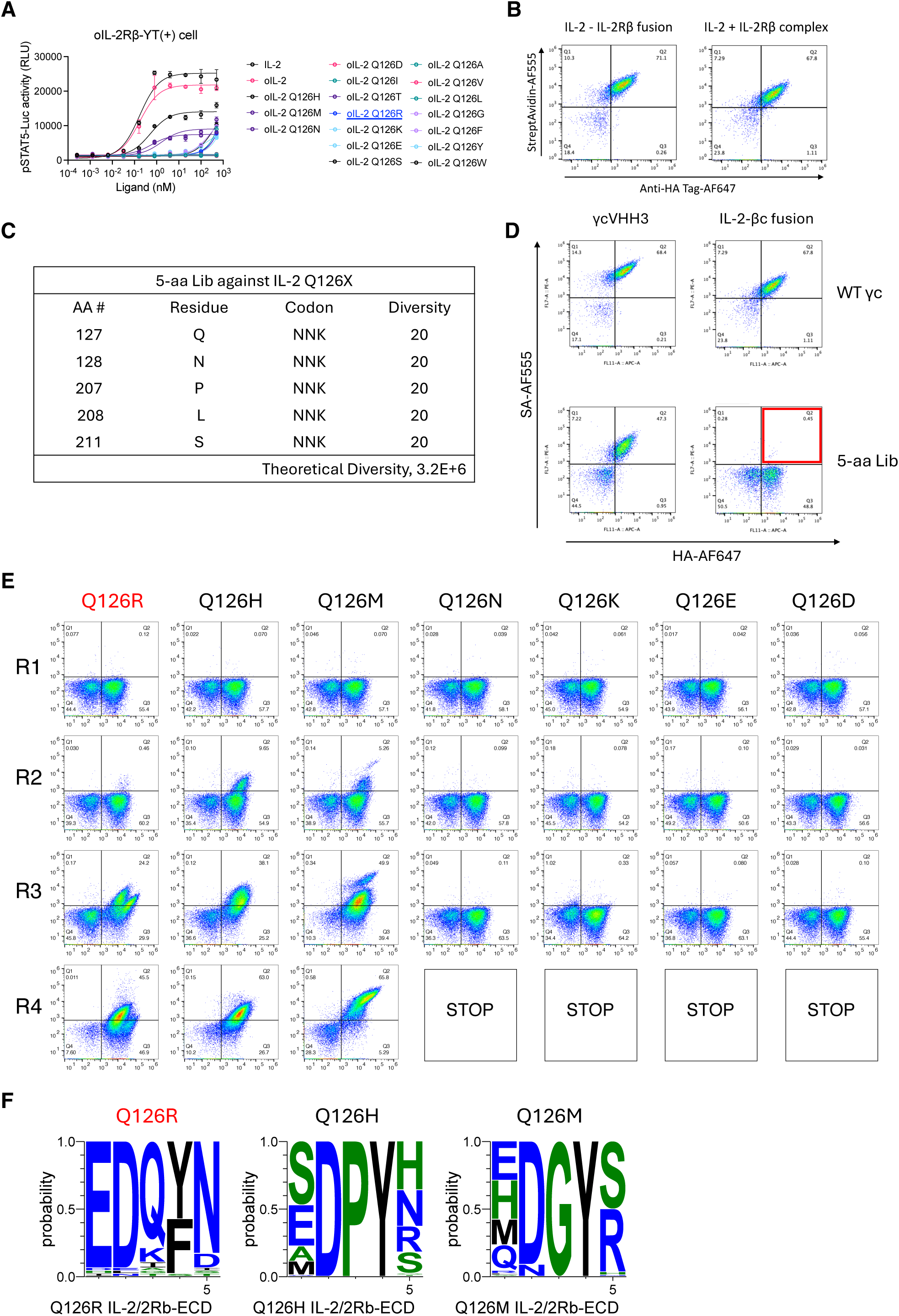
Directed evolution of orthogonal γc using yeast surface display, related to Figure 1. (A) Dose-response curves of pSTAT5 activity induced by serially diluted wild-type IL-2, single-orthoIL-2 (SoIL-2), and different double-orthoIL-2 (DoIL-2) candidates (SoIL-2 Q126X) in the STAT5-Luciferase YT+ reporter cell line expressing orthoIL-2Rβ and wild-type γc. Data were collected in duplicate and shown as mean ± SD. (B) Representative flow cytometry plots showing the staining of IL-2/IL-2Râ ECD protein complex and fusion against wild-type ãc on yeast surface. (C) Tables of mutated ãc amino acid positions and corresponding degenerate codons used to construct the mutant ãc yeast-display library. (D) Representative flow cytometry plots showing the yeast surface staining of wild-type ãc or mutant ãc library against reported ãc specific VHH and designed IL-2/IL-2Râ ECD fusion protein. (E) Representative flow cytometry plots showing the enrichment of the mutant ãc library during 4 rounds of evolution against different DoIL-2 candidates. (F) Sequence logos showing residue preferences of ãc variants after four rounds of yeast surface display selection against DoIL-2 candidates (Q126R, Q126N, and Q126M). Amino acid frequencies at each position are represented by letter size, with colors indicating residue identity.

**Figure S2.**
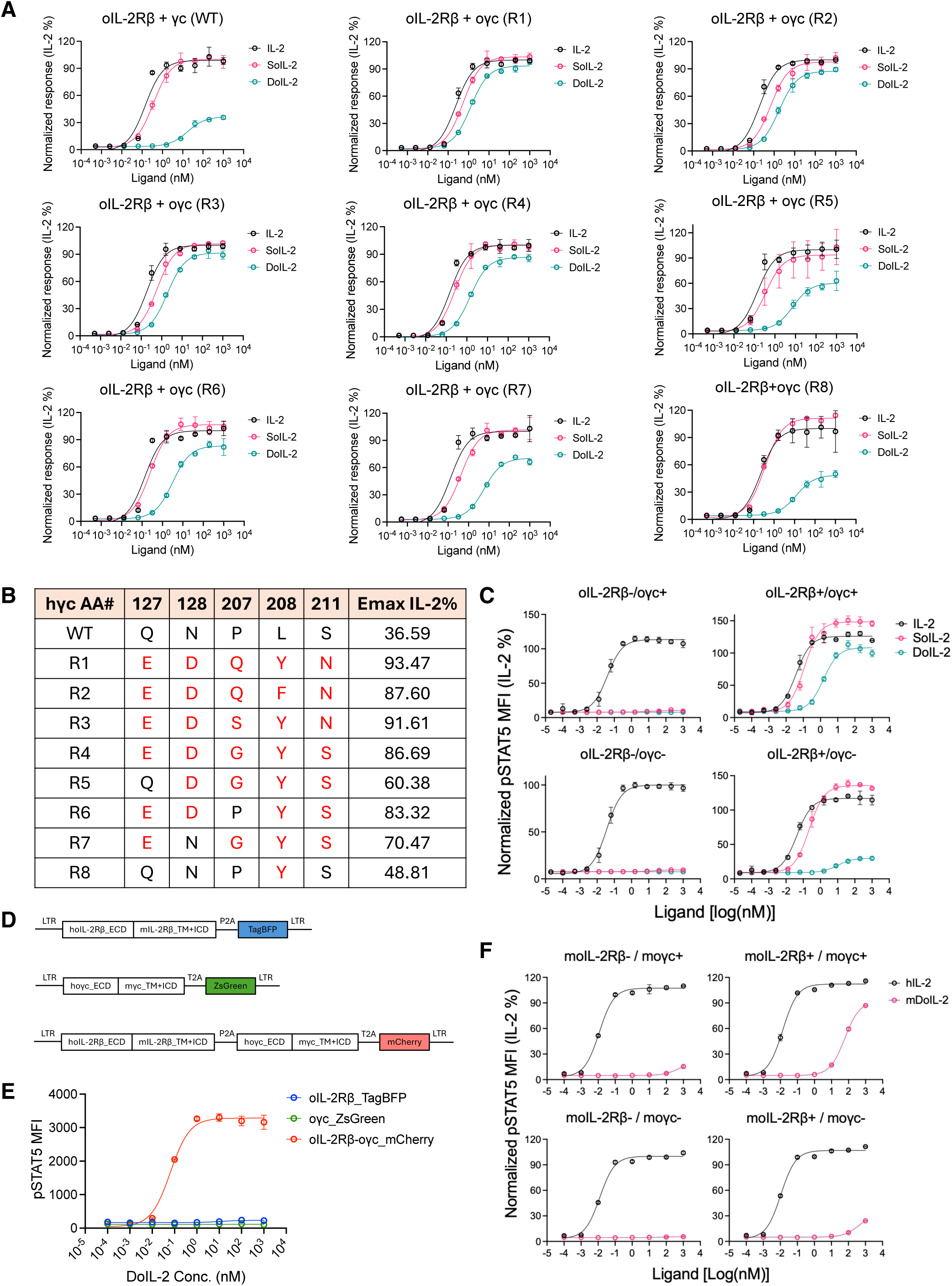
Functional validation of double-orthogonal IL-2/IL-2Rβ/γc signaling complex, related to Figure 1. (A) Dose-response curves of pSTAT5 activity induced by serially diluted wild-type IL-2, single-orthoIL-2 (SoIL-2), and double-orthoIL-2 (DoIL-2) in the STAT5-Luciferase YT+ reporter cell line expressing orthoIL-2Rβ and wild-type γc or different orthogonal γc candidates. Data were collected in duplicate and shown as mean ± SD. (B) Summary of amino acid sequences and relative pSTAT5 activity of all orthogonal γc candidates tested in (A). (C) Dose-response curves of STAT5 phosphorylation induced by IL-2, SoIL-2, or DoIL-2 in human primary T cells expressing ortho-IL2Râ and/or ortho-ãc. Median pSTAT5 MFI was normalized to wild-type IL-2-induced activity on double negative T cells. Data were collected in duplicate and shown as mean ± SD. (D) Construction of retroviral expression vectors for human-mouse chimeric ortho-IL-2Râ (TagBFP⁺), ortho-γc (ZsGreen⁺), and double-orthogonal receptors (mCherry⁺). A P2A peptide enabled coexpression of the two orthogonal receptors. (E) Dose-response curves of STAT5 phosphorylation induced by DoIL-2 in mouse primary T cells expressing human-mouse chimeric ortho-IL2Râ and/or ortho-ãc. Data were collected in duplicate and shown as mean ± SD. (F) Dose-response curves of STAT5 phosphorylation induced by human wild-type IL-2, or mouse DoIL-2 in mouse primary T cells expressing full-length mouse ortho-IL2Rβ and/or ortho-γc. Median pSTAT5 MFI was normalized to wild-type IL-2-induced activity on double negative T cells. Data were collected in duplicate and shown as mean ± SD.

**Figure S3.**
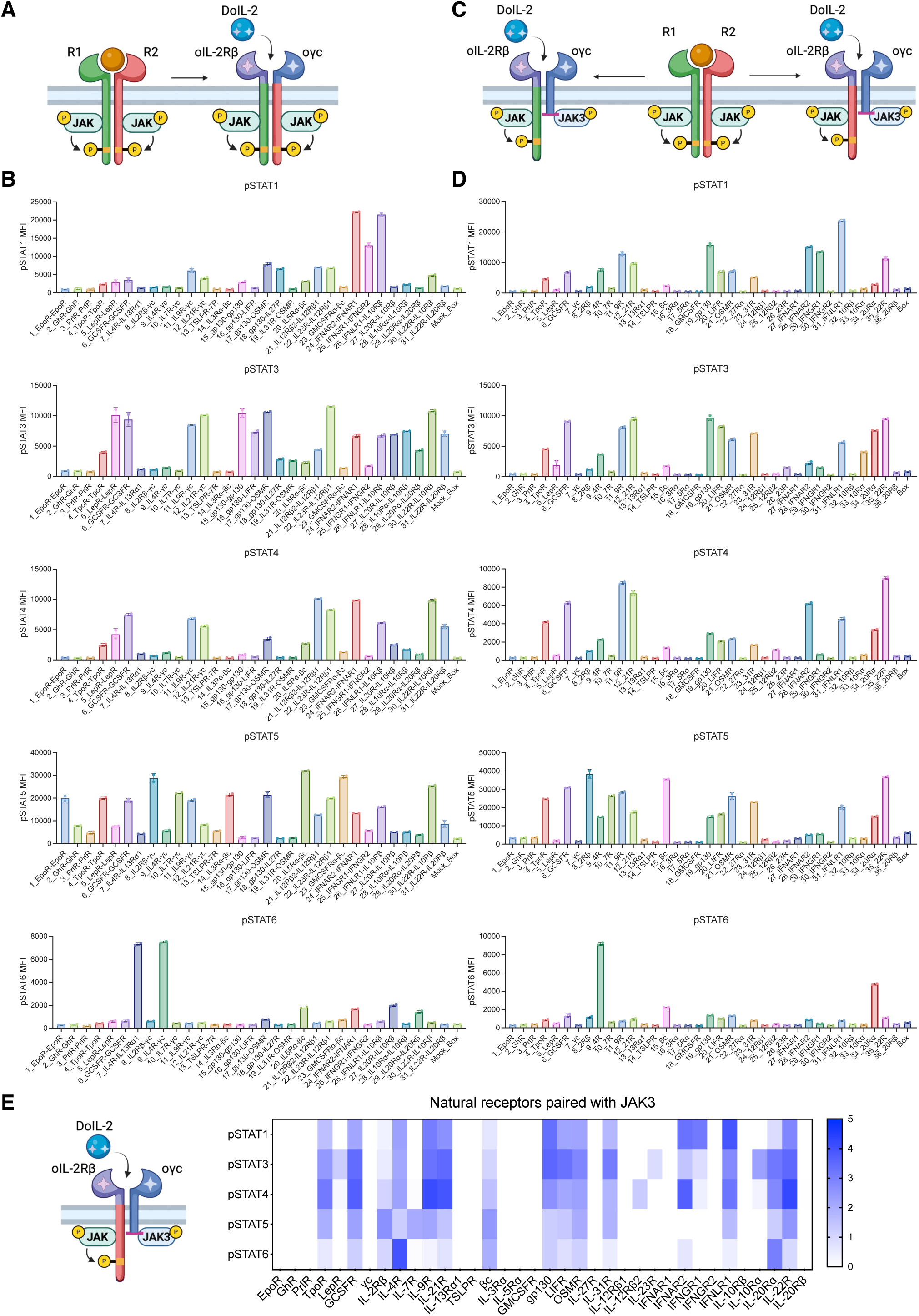
Construction and signaling profiles of dimeric and decoupled cytokine receptors, related to Figure 2. (A) Schematic of reconstructing dimeric cytokine receptor signaling from naturally occurring receptor pairs, achieved by swapping the intracellular domain of IL-2Rβ and γc. (B) Representative bar plots of raw median fluorescence intensity (MFI) for indicated pSTATs across reconstructed receptor pairs (n=31) from (A) after stimulation with 100 nM DoIL-2. Data were collected in duplicate and shown as mean ± SD. (C) Schematic of decoupling cytokine receptor signaling from naturally occurring receptor pairs, achieved by pairing a chimeric ortho-IL-2Rβ receptor with an ortho-γc containing the only JAK-binding region. (D) Representative bar plots of raw median fluorescence intensity (MFI) for indicated pSTATs across decoupled receptors (n=36) from (C) after stimulation with 100 nM DoIL-2. Data were collected in duplicate and shown as mean ± SD. (E) Heatmap of relative pSTAT1/3/4/5/6 signaling activity across all decoupled receptors (n=36) described in (C and D).

**Figure S4.**
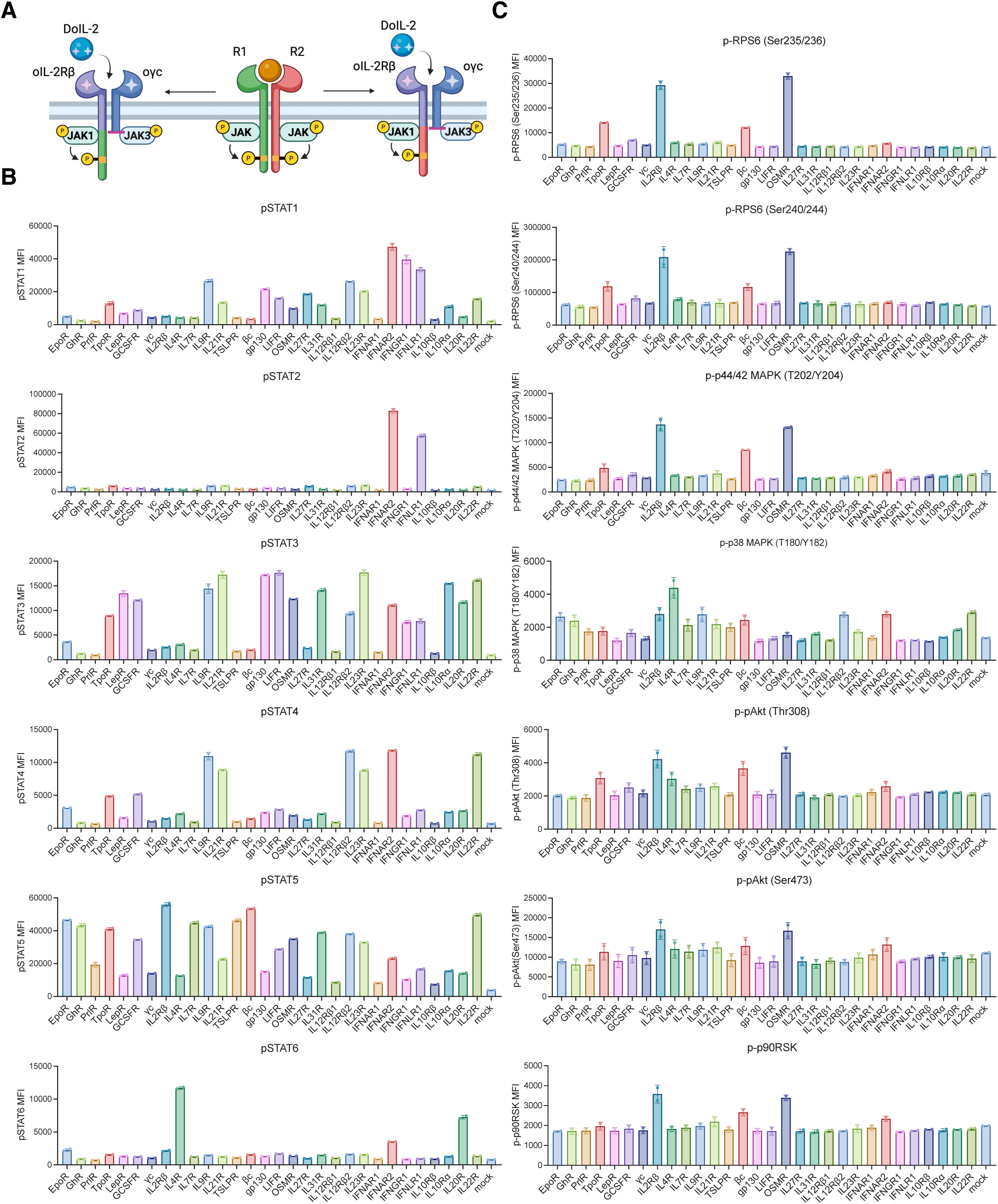
Construction and signaling profiles of decoupled cytokine receptors with fixed JAK-binding region, related to Figure 2. (A) Schematic of decoupling cytokine receptor signaling from naturally occurring receptor pairs, achieved by pairing an optimized chimeric ortho-IL-2Rβ receptor harboring fixed JAK-binding region with an ortho-γc containing the only JAK-binding region. (B) Representative bar plots of raw median fluorescence intensity (MFI) for indicated pSTATs across decoupled receptors (n=30) from (A) after stimulation with 100 nM DoIL-2. Data were collected in duplicate and shown as mean ± SD. Mouse STAT2 activity was measured using transduced human STAT2 as a surrogate readout due to lack of suitable detection antibodies. (C) Representative bar plots of raw median fluorescence intensity (MFI) for phosphorylation of non-STAT signaling molecules across decoupled receptors (n = 30) defined in (A) following stimulation with 100 nM DoIL-2. Data were collected in duplicate and shown as mean ± SD.

**Figure S5.**
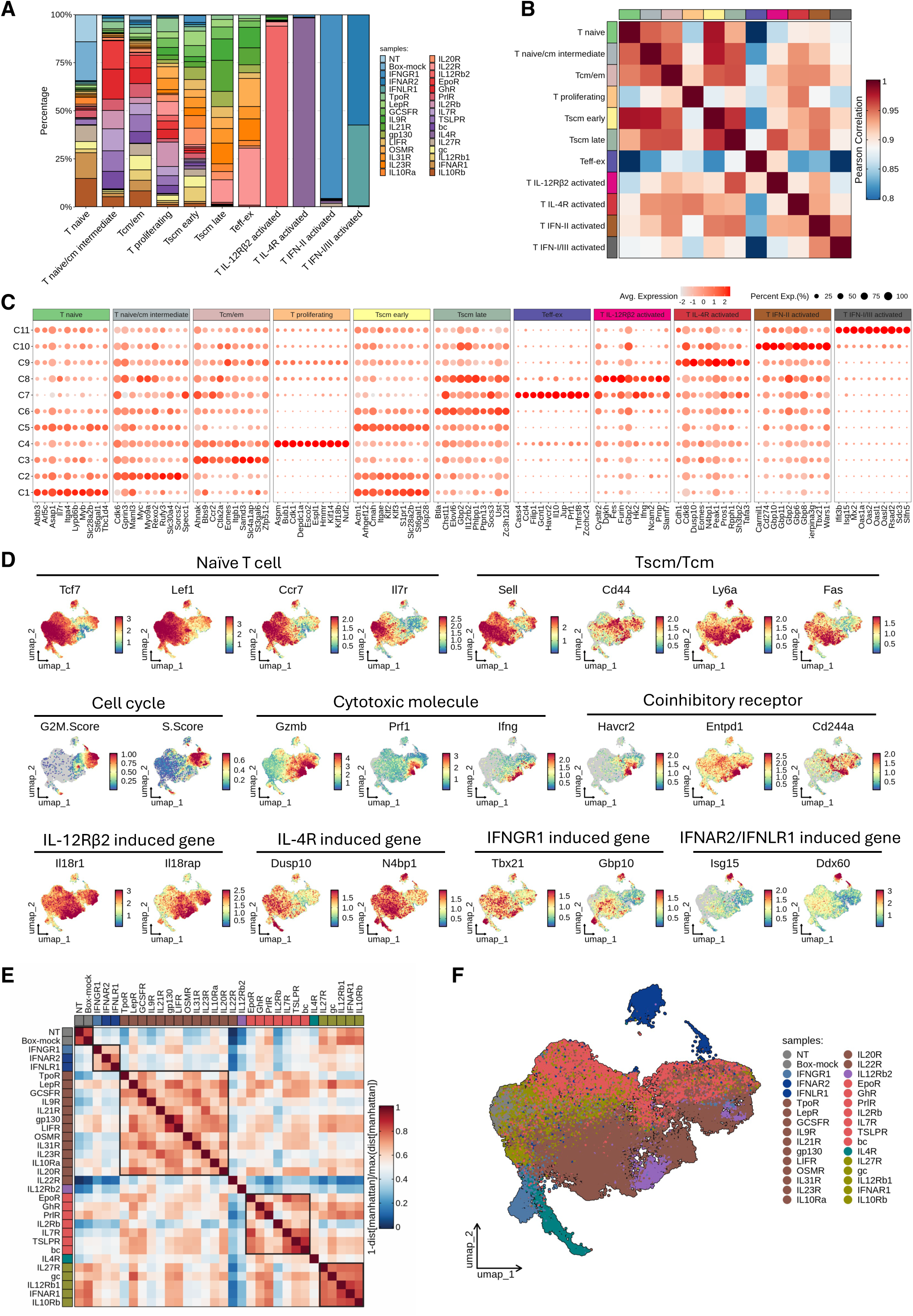
Characterization of T cell transcriptional signatures across all orthogonal cytokine receptor signaling conditions, related to Figure 3. (A) Stacked bar plot showing the proportion of cells across samples for each annotated cluster, colored by sample condition. (B) Heatmap of Pearson correlation coefficients across annotated T cell clusters based on transcriptomic profiles. Each cell represents the pairwise correlation between clusters, with color indicating correlation strength. (C) Bubble heatmap showing expression of top marker genes for each cluster identified by differential expression analysis. Dot size represents the percentage of cells expressing each gene, and color intensity indicates normalized expression levels. (D) UMAP feature plots showing expression of representative genes associated with major T cell states and functional programs, including naïve, Tscm/Tcm, cell cycle, cytotoxicity, coinhibitory receptors, and specific cytokine receptor-induced gene signatures. Color indicates normalized gene expression levels. (E) Heatmap of pairwise distances between samples across all cytokine receptor conditions based on transcriptomic profiles. Distances were calculated using the Manhattan metric and transformed as 1 − distance/max(distance). (F) UMAP visualization of scRNA-seq data from all collected T cells, colored by cytokine receptor conditions, showing the distribution and overlap of cells across conditions.

**Figure S6.**
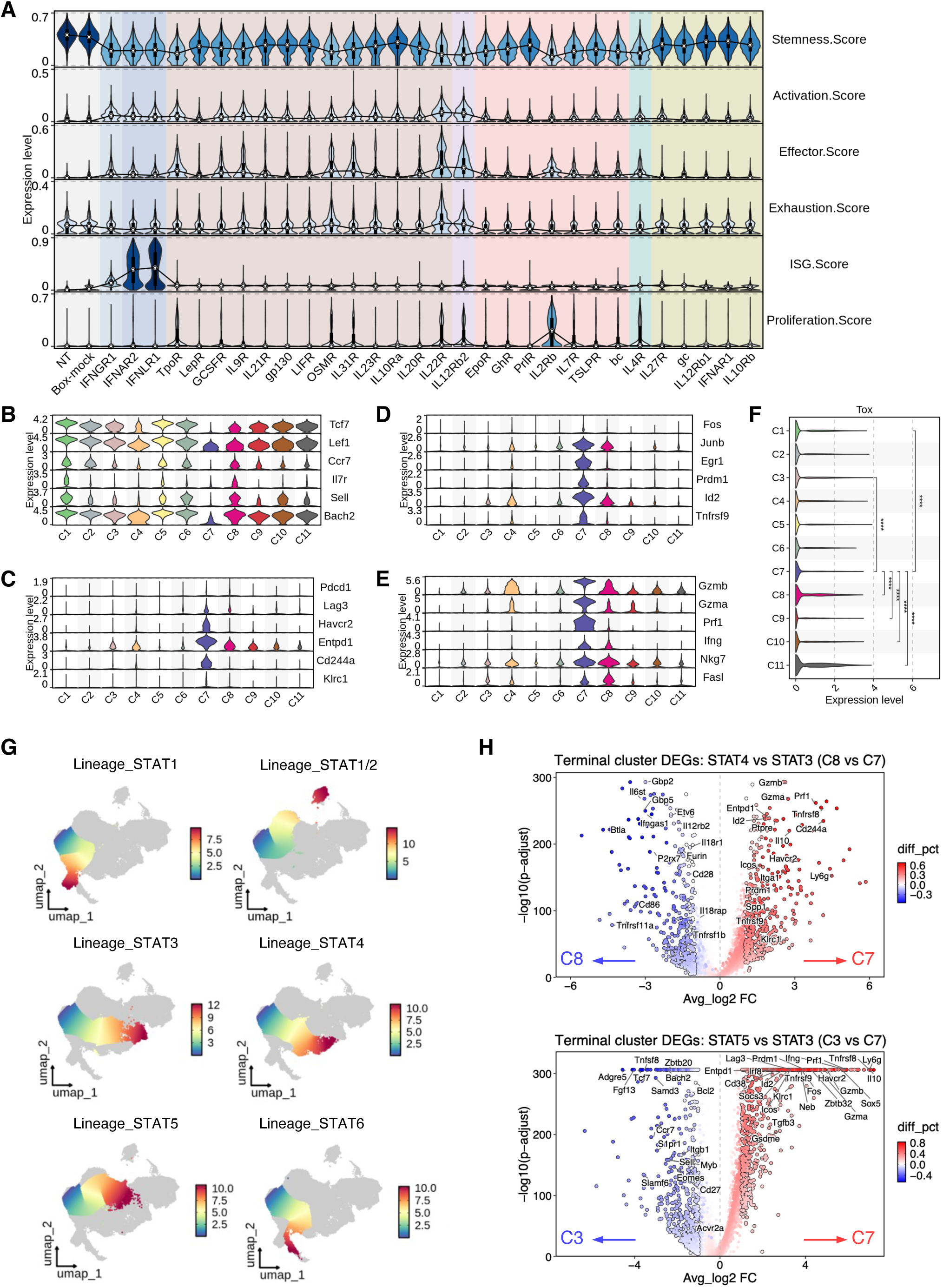
Functionally divergent T cell differentiation programs driven by distinct STAT signaling downstream of cytokine receptors, related to Figure 4. (Α) Stacked violin plots showing relative expression of the signature score for stemness, activation, effector, exhaustion, ISG, and proliferation across all cytokine receptor conditions. (Β-Ε) Violin plots showing relative expression of representative genes from the gene sets used to calculate the signature score for stemness (Β), exhaustion (C), activation (D), and effector (E) across annotated T cell clusters. (F) Differential expression analysis of Tox across annotated T cell clusters. Violin plots showing relative expression levels with statistical significance indicated. (G) UMAP visualization of six Slingshot-inferred lineages of non-cycling T cells corresponding to distinct STAT programs. (H) Differential gene expression between the STAT3-driven terminal cluster (C7) and the STAT4-(C8) or STAT5-driven (C3) terminal clusters. Volcano plots highlight genes enriched in distinct STAT-driven T cell differentiation clusters, with color denoting the difference in the fraction of expressing cells (diff_pct).

**Figure S7.**
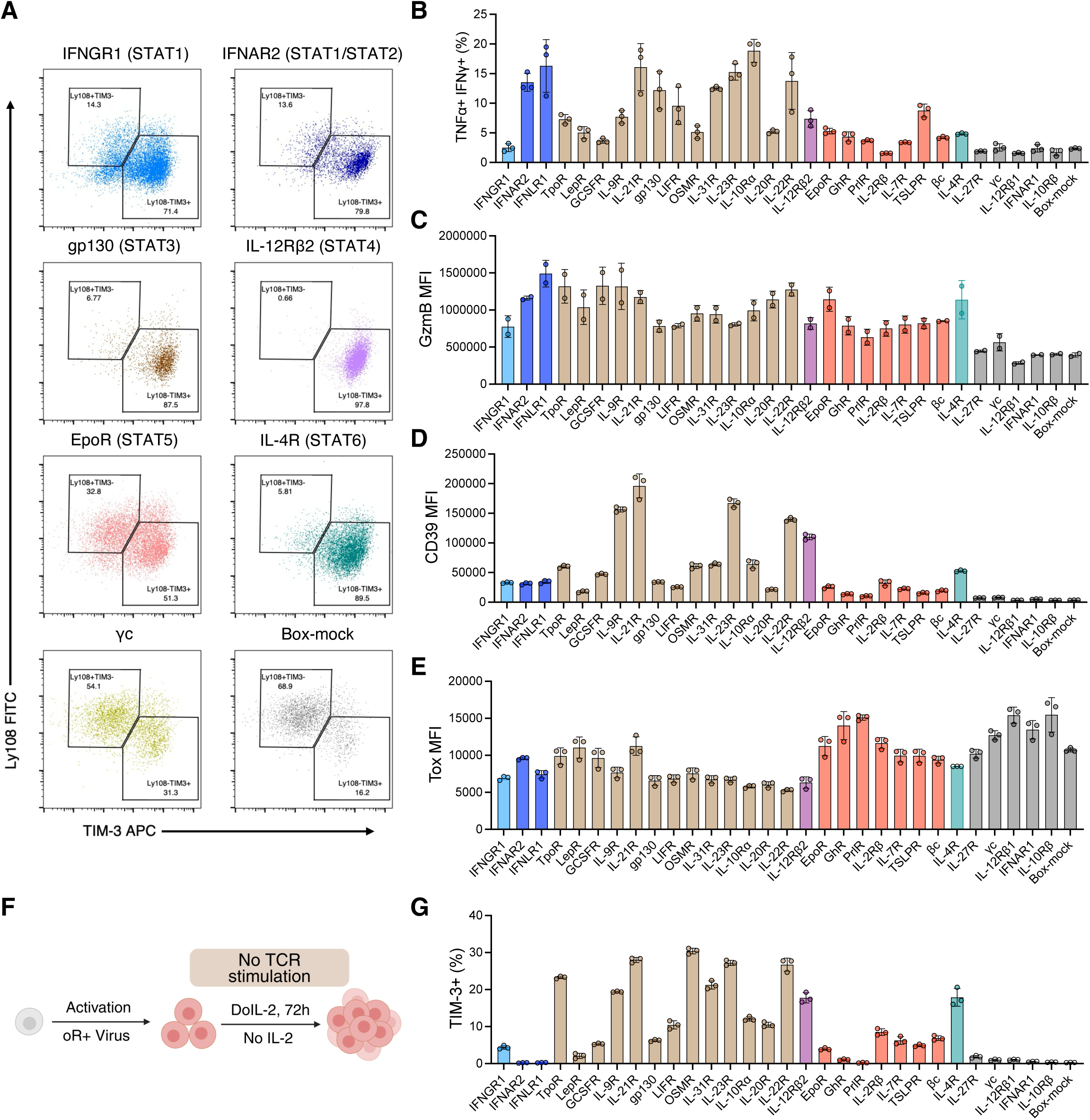
Functional and phenotypic characterization of cytokine receptor-programmed T cells, related to Figure 4. (A) Representative flow cytometry plots of TIM-3 and Ly108 expression within the CD44^+^PD1^+^ gate in T cells stimulated through indicated STAT-specific cytokine receptors, distinguishing progenitor (Tex_prog; Ly108⁺ TIM-3⁻) and terminal (Tex_term; Ly108⁻TIM-3⁺) exhausted T cell populations. (B) TNFá^+^IFN-γ^+^ cytokine-producing T cell frequencies after repeated CD3/CD28 stimulation across all receptor conditions, ordered by dominant STAT category. Data represent three biological replicates, mean ± SD. (C-E) Bar plots showing the differential expression level of GzmB (C), CD39 (D) and Tox (E) of T cells after repeated CD3/CD28 stimulation across all cytokine receptor conditions as shown in Figure 4E. Data are representative of three biological replicates and shown as mean ± SD. (F) Schematic of the *in vitro* 3-day DoIL-2 stimulation of T cells without TCR activation. (G) Bar plots showing the differential frequencies of TIM-3^+^ T cells following 3 days of *in vitro* DoIL-2 stimulation across all cytokine receptor conditions in the absence of CD3/CD28 stimulation. Data are representative of three biological replicates and shown as mean ± SD.

**Figure S8.**
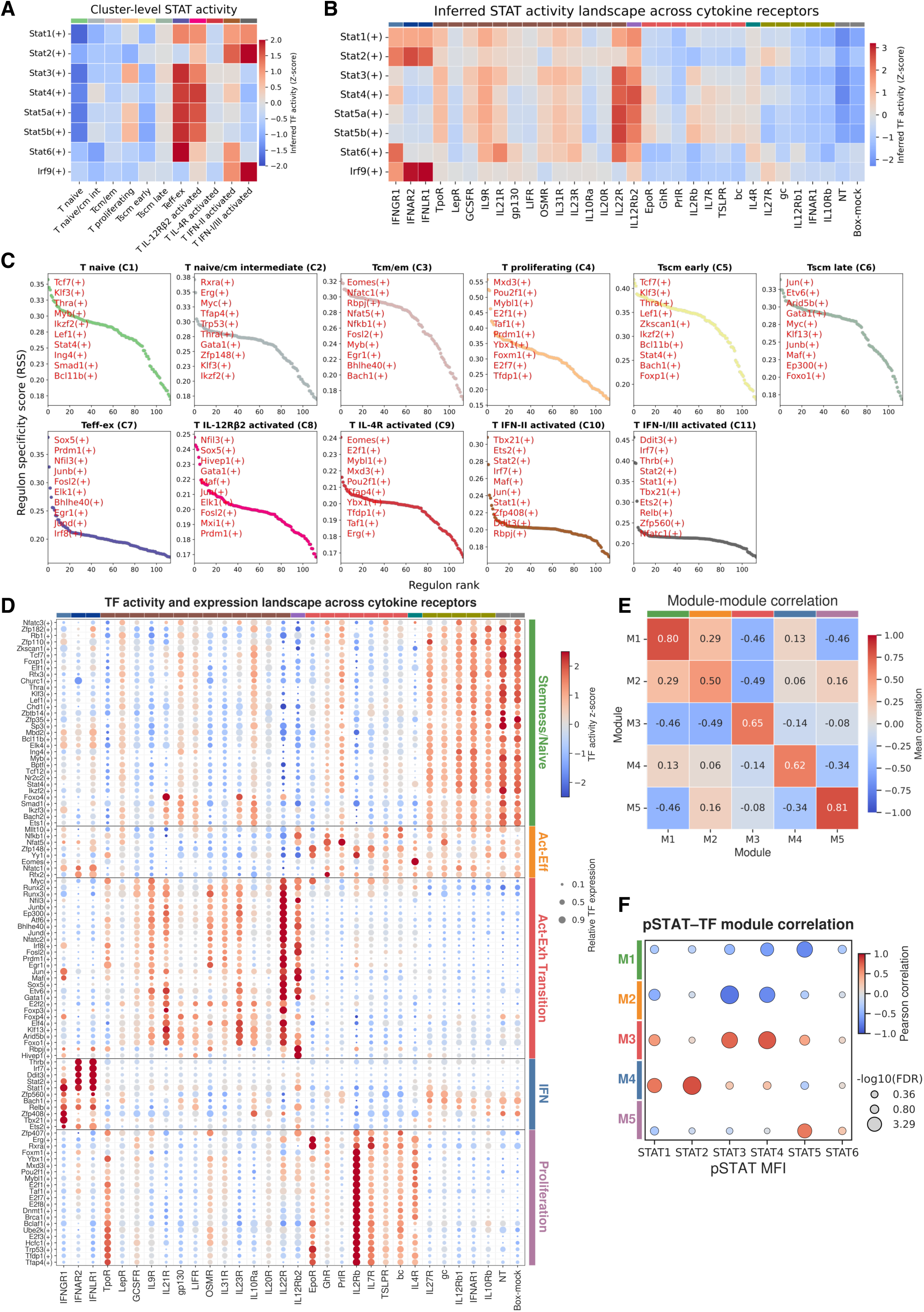
Inferred transcription factor activity in T cells across annotated clusters and cytokine receptor conditions, related to Figure 5. (A) Heatmap of cluster-level STATs and IRF9 activity inferred using ULM with CollecTRI regulons. Color denotes scaled TF activity (z-score). (B) Heatmap of receptor-level STATs and IRF9 activity inferred using ULM with CollecTRI regulons. Color denotes scaled TF activity (z-score). (C) Regulon specificity score (RSS) ranking across annotated T cell clusters. Regulons are ordered by cluster specificity, with top 10 cluster-specific transcription factors highlighted in red. (D) Bubble heatmap showing SCENIC-inferred regulon activity and corresponding gene expression across all cytokine receptor conditions. Regulons are classified into five indicated modules. Color denotes scaled regulon activity (AUC), and dot size represents relative gene expression. (E) Heatmap showing mean pairwise correlation between regulatory modules identified by hierarchical clustering. Each value represents the mean pairwise regulon correlations between modules. (F) Correlation between pSTAT MFI and TF regulatory module activity across all receptor conditions. Color indicates correlation direction and strength; dot size reflects statistical significance. Each receptor’s STAT activation profile predicts its downstream TF regulatory program.

**Figure S9.**
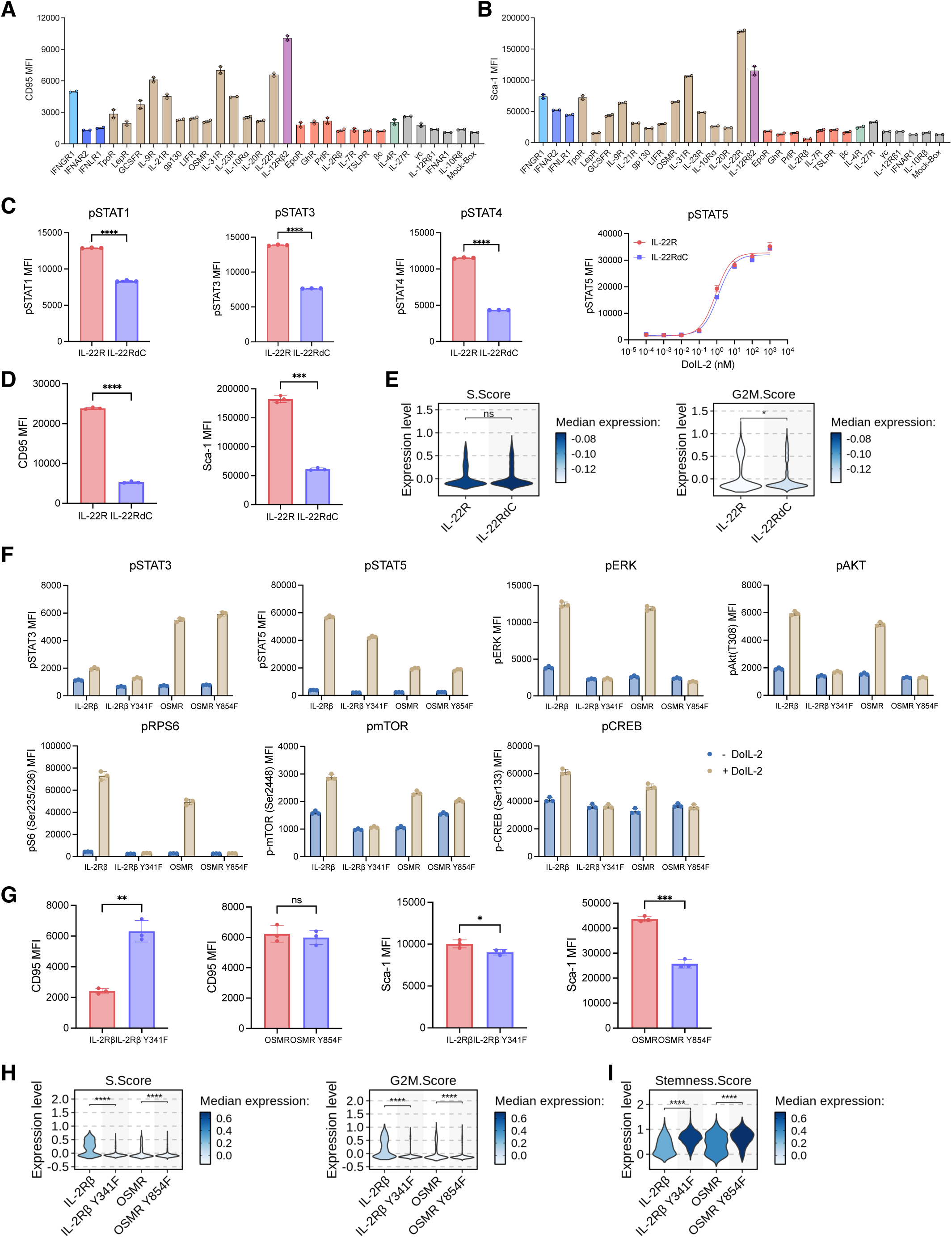
Engineering cytokine receptor intracellular domain to rewire downstream signaling rebalances T cell proliferation and stemness, related to Figure 6. (A-B) Bar plots showing the differential expression level of Tscm-related markers Fas/CD95 (A) and Sca-1 (B) following 3 days of stimulation across all orthogonal cytokine receptor conditions as shown in Figure 6B. Data are representative of biological duplicates and shown as mean ± SD. (C) Characterization of STAT signaling in T cells expressing orthogonal IL-22R or IL-22RdC. (D) Comparison of CD95 and Sca-1 expression levels in T cells expressing orthogonal IL-22R or IL-22RdC following *in vitro* stimulation. (E) Violin plots showing S phase and G2/M phase scores in T cells expressing orthogonal IL-22R or IL-22RdC. Comparisons were derived from scRNA-seq data. Statistical significance is indicated. (F) Characterization of STAT and Non-STAT signaling in T cells expressing orthogonal IL-2Râ or its Y341F variant, and OSMR or its Y854F variant. (G) Comparison of CD95 and Sca-1 expression levels in T cells expressing orthogonal IL-2Rβ or its Y341F variant, and OSMR or its Y854F variant following *in vitro* stimulation. (H-I) Violin plots showing S phase and G2/M phase scores (H) and stemness score (I) in T cells expressing orthogonal IL-2Rβ or its Y341F variant, and OSMR or its Y854F variant. Comparisons were derived from scRNA-seq data. Statistical significance is indicated.

**Figure S10.**
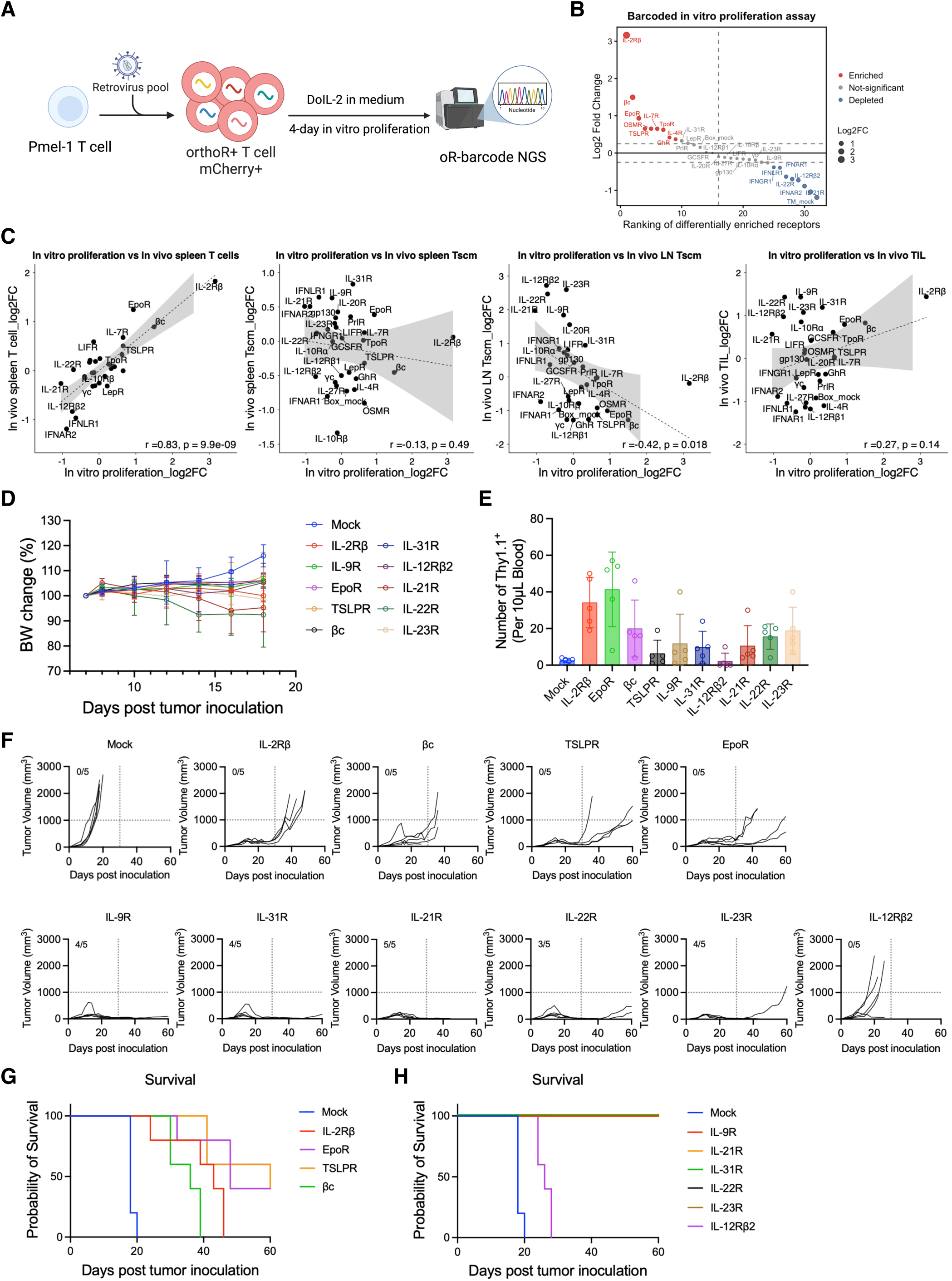
*In vivo* profiling of antitumor efficacy across selected orthogonal cytokine receptors in the B16F10 model, related to Figure 7. (A) Schematic of *in vitro* selection of enriched T cells expressing barcoded receptors following DoIL-2 stimulation. (B) Scatter plot showing differential enrichment of barcoded cytokine receptors following *in vitro* selection. Receptors are ranked by barcode enrichment, with log2 fold change shown. Color denotes enriched, depleted, or not significant receptors, and dot size reflects effect size. (C) Correlation analysis showing the relationship between receptor enrichment *in vitro* and *in vivo*, across T cell subsets as shown in Figure 7D. (D) Relative body weight change in mice following the indicated ACT treatments. (E) Number of transduced Pmel-1 CD8⁺ T cells in 10 μL of tail blood 5 days after the indicated ACT treatments. (F) Individual B16F10 tumor growth curves of C57BL/6 mice treated with Pmel-1 T cells transduced with indicated orthogonal cytokine receptors. The number of tumor-free mice relative to the total number per group is indicated. (G) Survival curves of mice treated with Pmel-1 T cells expressing orthogonal cytokine receptors with dominant STAT5 signaling. (H) Survival curves of mice treated with Pmel-1 T cells expressing orthogonal cytokine receptors with dominant STAT3 or STAT4 signaling.

